# Enhancing Neural Synchrony with Endogenous-like 1/f Noise Stimulation

**DOI:** 10.1101/2025.04.30.651370

**Authors:** Anan Ping, Longzhou Guan, Hai Gu, Zhipeng Jiang, Wenhao Deng, He Chen, Ke Zeng, Xiaoli Li

**Author notes:** Anan Ping, Longzhou Guan and Hai Gu contributed equally to this work. Correspondence to: He Chen, Ke Zeng, Xiaoli Li.

## Abstract

Aperiodic components of neural activity, characterized by endogenous 1/f noise dynamics, are hypothesized to support the emergence of large-scale cortical order and cognitive flexibility. Here, we combine computational modeling and human brain stimulation to elucidate the role of 1/f noise in modulating neural synchrony. Using a coupled oscillator model, we demonstrate that ubiquitous 1/f noise does more effectively enhances phase synchrony than spectrally flat (white) noise. Crucially, we identify a competitive synergy between noise intensity and the 1/f spectral exponent: starting from optimal white noise-induced synchrony, increasing the 1/f exponent while decreasing noise intensity leads to a further enhancement of synchrony, which peaks at a specific parameter regime before diminishing. To experimentally validate these findings, we developed a transcranial 1/f noise stimulation (tFNS) system and applied it to human subjects. Compared to spectrally white noise stimulation, the tFNS more robustly enhanced corticospinal synchrony, consistent with model predictions. These results uncover a functional advantage of scale-free brain noise in driving coordinated neural dynamics, offering a new framework for optimizing non-invasive brain stimulation. More broadly, our findings suggest that the brain may harness stochastic facilitation through adaptive modulation of its aperiodic activity to support ordered macro-dynamics.

## Introduction

The human brain operates as a spontaneously noisy nonlinear system^1^, characterized by complex, chaotic firing patterns across neuronal networks and pervasive electrochemical fluctuations^2, 3^. These networks function as coupled oscillators^4, 5^, and their aggregate electrical activity exhibits a power-law distribution in the power spectral density^6, 7^—commonly described as 1/f scaling, or the aperiodic component of neural activity^8^. This component, parameterized by the 1/f exponent, reflects the non-periodic background signal (Fig. 1a), which was previously regarded as the harmful noise part^9^, and has recently emerged as a critical substrate for the hierarchical coordination of neural dynamics across multiple spatial and temporal scales^10^.

**Figure 1.**
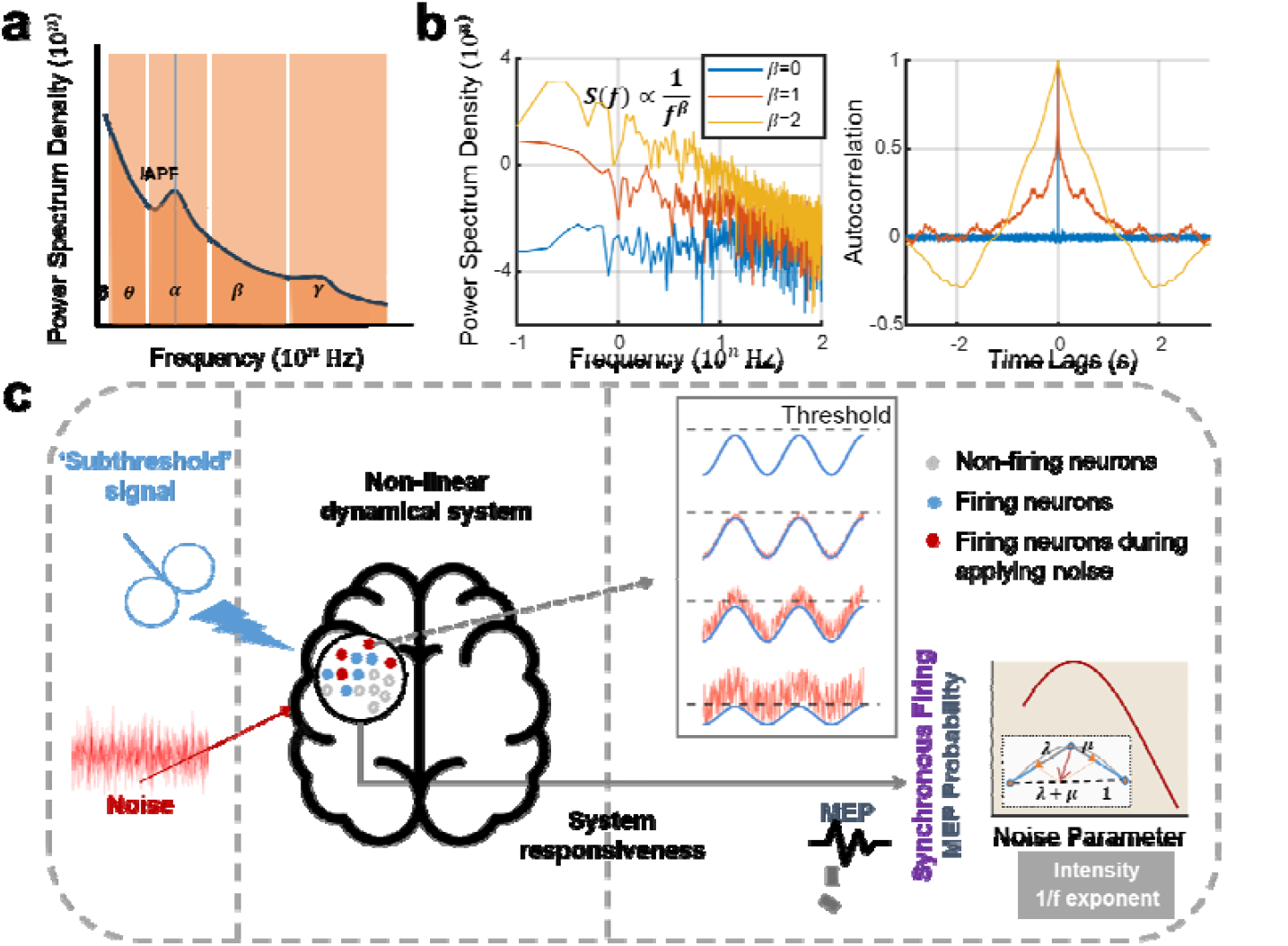
Schematic illustration of stochastic facilitation and biologically relevant noise in neural systems. **a**. Power spectral density of spontaneous brain electrical activity displays a characteristic 1/f structure, indicative of scale-free (aperiodic) dynamics in neural systems. **b**. Comparison of noise types. Unlike white noise (1/f exponent *β* = 0, blue), which has no temporal correlation and a flat frequency spectrum, 1/f-like noise exhibits long-range temporal correlations and power-law decay in the frequency domain. Pink noise (*β* = 1, red) and brown noise (*β* = 2, yellow) show increasingly steep slopes, corresponding to stronger temporal correlations and greater low-frequency dominance. **c**. Conceptual framework and hypothesis. A subthreshold signal (analogous to weak TMS) does not elicit a detectable response in the absence of noise, but appropriately tuned 1/f noise enhances system responsiveness (synchronous firing of corticospinal neurons, the macroscopic manifestation is MEPs) via stochastic facilitation. Excessive noise, by contrast, degrades signal transmission. A convexity index (CI) is proposed to quantify the characteristic inverted-U relationship between system responsiveness (MEP probability in TMS-MEP experiments and synchronization index in Kuramoto model) and noise parameters (intensity and 1/f exponent).

Accumulating evidence suggests that the 1/f component underlies key features of brain activity, including the long-tail distribution of neuronal avalanches and the formation of large-scale organized patterns^7, 11^. Such scale-free dynamics may enable the brain to sustain a flexible dynamical repertoire and rapidly adapt to changing stimuli^12, 13^. Alterations in the 1/f structure have been associated with disruptions in excitation–inhibition balance^14^, abnormalities in consciousness and age-related neural decline^15^. Given its widespread and conserved presence across cortical systems^7, 13, 16^, 1/f noise represents a biologically meaningful class of input that mimics endogenous brain fluctuations. Unlike white noise, which distributes energy equally across frequencies, 1/f noise exhibits an inverse relationship between power and frequency (Fig. 1b), prompting the question of whether endogenous-like 1/f noise input could more effectively enhance neural synchrony via stochastic facilitation (SF)^7^.

Phenomenon that appropriate noise facilitating nonlinear system responsiveness^17^, is commonly referred to as stochastic facilitation (SF) in neuroscience^7^. Growing evidence confirmed that brain’s information processing can benefit from added noise^3, 18, 19^: applying transcranial random noise stimulation (tRNS) on visual cortex could improve the detection of low-contrast visual stimuli^20^, decision-making^21^, visual-motor perception^22^, visual training and boost visual recovery in cortically blind patients^23^. This benefit was also found at the action potential of single neurons^24, 25^ in vitro and cortical neuronal excitability^26^. However, despite the increased interest about SF in human brain, most researches have been limited to white noise^27^, and the variability in outcomes across tRNS studies may reflect unresolved uncertainties in noise delivery protocols^20, 28^. Hence, one of the most interesting unresolved scientific question about SF in neuronal systems is to explore the biologically relevant noise and understand how the relevant noise plays a role in human brain. Here, we hypothesize that the brain preferentially benefits from 1/f noise over spectrally flat noise through enhanced stochastic facilitation, and that this benefit depends on an interaction between noise intensity and the 1/f exponent.

To test this, we employed numerical simulations based on the Kuramoto oscillator model— capturing phase synchronization among coupled neural populations—and validated the predictions experimentally using transcranial magnetic stimulation–evoked motor potentials (TMS-MEP), a physiological index of corticospinal neuron synchrony. We introduced a convexity index (CI) to quantify the hypothesized non-monotonic (inverted U-shaped) relationship between synchronization and noise parameters (Fig. 1c). Our findings reveal a synergistic interaction between the intensity and spectral structure of noise in modulating neural synchrony. They underscore the functional relevance of the brain’s endogenous aperiodic dynamics and provide the first empirical demonstration that transcranial 1/f noise stimulation (tFNS) can modulate subthreshold signal processing. These insights advance our understanding of stochastic facilitation in neural systems and establish a principled framework for optimizing biologically inspired brain stimulation protocols.

## Result

### Oscillator Synchronization Exhibits an Inverted U-Shaped Dependence on White Noise Intensity in the Kuramoto Model

To examine the influence of noise on neural synchronization, we employed the Kuramoto model, a paradigmatic framework for simulating coupled phase oscillators. The coupling strength between oscillators is governed by a parameter *k*, where *k* = 0 results in no interaction (i.e., no phase synchrony, *R* = 0), and sufficiently large *k* leads to full synchronization. The critical value at which synchronization first emerges is denoted *k*_critical_ = 1. Accordingly, we define coupling regimes as *subthreshold* (*k* < 1) and *suprathreshold* (*k* > 1). To systematically explore the interaction between noise and coupling strength, we conducted simulations across a two-dimensional parameter space: *k* was varied from 0 to 1.5 in steps of 0.05, and white noise intensity *σ* from 0 to 4 in steps of 0.1 (Model 1). The goal was to determine how noise intensity modulates the synchronization index *R*, and whether an optimal value of *σ* exists that maximizes synchronization.

As shown in Figure 2a, in the near-subthreshold range of coupling strengths k (orange shaded area), the synchronization index *R* is markedly higher under moderate white noise intensities (*σ* = 1 and 2) compared to the no-noise condition (*σ* = 0). This indicates that white noise can enhance phase synchronization when the system operates just below the critical coupling threshold. In contrast, for ultra-low coupling strengths (*k* is close to 0, green circle), noise fails to reliably induce synchronization, and in the suprathreshold regime (*k* > 1, yellow circle), increasing noise disrupts the already established synchrony.

**Figure 2.**
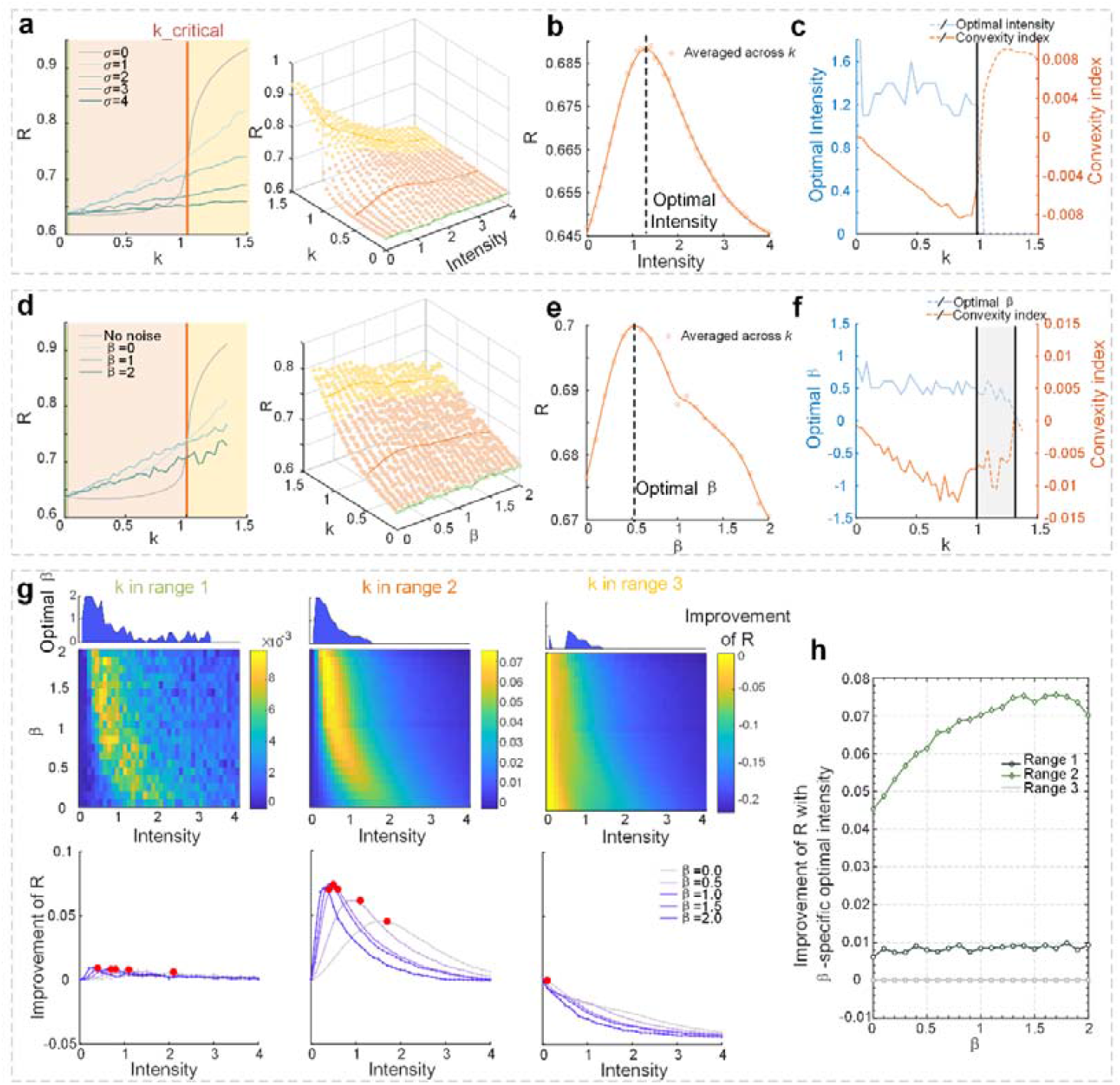
Synergistic effects of noise intensity and 1/f exponent on oscillator synchronization. **a**. Synchronization index R as a function of coupling strength k, shown for representative white noise intensities *σ* = 0,1,2,3,4. When *σ* = 0, there is a sigmoid function. **b**. Surface plot of R across the (*k,σ*) parameter space. Coupling strength *k* varies from 0 to 1.5; noise intensity *σ* varies from 0 to 4 (step size = 0.1). Three coupling regimes are indicated: ultra-low *(k* < 0.1, green: no enhancement), near-subthreshold (0.1 < *k* < 1, orange: noise-enhancing), and suprathreshold (*k* > 1, yellow: noise-disrupting). **c**. Inverted U-shaped relationship between *R* and *σ*, averaged across the near-subthreshold coupling range. **d**. Optimal noise intensity *σ* (blue) and convexity index (CI, red) as functions of *k*. CI < 0 indicates convex (inverted-U) behavior; the zero-crossing of CI corresponds to the synchronization emergence threshold at *k*=1. **e–f**. Same analysis framework as in a–d, but using 1/*f* noise with fixed intensity *σ* = 1 and variable 1/f exponent *β*∈[0,2]. **g**. Top panel: heatmap of synchronization improvement (relative to the no-noise baseline) as a function of *σ* and *β*, averaged over three *k* regimes: ultra-low, near-subthreshold, and suprathreshold (defined via CI in Supplementary Fig. S16). Bottom panel: cross-sectional curves of synchronization improvement versus *σ* for representative values of *β* = 0, 0.5, 1.0, 1.5, 2.0 across the three *k* regimes. Red circles mark the local maxima, denoting optimal *σ* values for each *β*. **h**. Synchronization improvement as a function of *β*, where *σ* is fixed at the optimal value for each *β*, shown separately for the three *k* regimes.

Moreover, when *k* lies in the subthreshold regime, the relationship between synchronization *R* and noise intensity *σ* exhibits an inverted U-shaped profile (Figure 2a, right panel, orange circle), suggesting the existence of an optimal noise level that maximally promotes synchronization. Averaging across the near-subthreshold values of *k*, we found that the synchronization peaked at *R*_max_ = 0.69 when *σ*_optimal_ = 1.4 (Figure 2b).

To quantitatively capture this non-monotonic behavior, we introduced a convexity index (CI) (see Materials and Methods). A negative CI indicates an inverted-U trend, with larger absolute values reflecting stronger convexity. As shown in Figure 2c, both the optimal noise intensity and CI are plotted across the full range of coupling strengths k. Notably, the critical transition point—where CI crosses zero—coincides precisely with the synchronization emergence threshold at *k* = 1. In the ultra-low range (*k* < 0.1), CI values remain positive, indicating a lack of convexity and thus no optimal enhancement. Therefore, the CI analysis confirms that optimal noise-induced synchronization is confined to the subthreshold interval (0.1 < *k* < 1), and is absent in both the suprathreshold and ultra-low regimes.

### 1/f Noise Enhances Oscillator Synchronization More Effectively Than White Noise in the Kuramoto Model

1/f noise is characterized by two key parameters: the noise intensity *σ* and the 1/f spectral exponent *β*. To isolate the influence of the 1/f exponent on synchronization, we fixed the noise intensity at *σ* = 1 and systematically varied *β* from 0 to 2 in increments of 0.1, across coupling strengths *k*∈[0, 4/3] in steps of 0.1 (Model 2). As shown in Figure 2d, for near-subthreshold coupling strengths (0.1 < *k* < 1), all noise conditions (*β* = 0,1,2) improved synchronization *R* relative to the no-noise baseline. In contrast, neither the ultra-low (*k* < 0.1) nor suprathreshold (*k* > 1) regimes exhibited consistent noise-enhancing effects. Notably, 1/*f* noise with higher spectral exponents (*β* = 1,2) yielded stronger enhancement of *R* than white noise (*β* = 0).

In the right panel of Figure 2d, an inverted U-shaped relationship between synchronization *R* and *β* is observed when *k* is in the subthreshold range, suggesting the existence of an optimal 1/f exponent. Averaging across the subthreshold *k* values yielded a peak synchronization of *R*_*max*_ = 0.7 at *β*_optimal_ = 0.5 (Figure 2e). We quantified this non-monotonic relationship using the convexity index (CI), with results shown in Figure 2f. Interestingly, CI remained negative up to *k* = 1.3, indicating that optimal *β* values—and associated enhancements in synchronization—persist slightly beyond the critical coupling threshold. This suggests that 1/f noise not only provides superior enhancement compared to white noise in the subthreshold regime but also imposes less destructive interference in weakly suprathreshold conditions.

### Optimal Balance between Noise Intensity and 1/f Exponent Maximizes Synchronization in the Kuramoto Model

To explore the interaction between noise intensity *σ* and 1/*f* exponent *β* in modulating synchronization, we performed simulations across a two-dimensional grid with *β*∈[0,2] and *σ*∈ [0,4], each in increments of 0.1. Given that the effect of noise depends on coupling strength, we stratified the analysis into three intervals based on convexity index values (see *Supplementary Materials* Fig. S16): Ultra-low coupling: k∈[0,0.1], CI > 0; Near-subthreshold: k∈[0.1,1], CI < 0;

Suprathreshold: k∈[1,4/3], CI > 0. For each interval, synchronization values were averaged across *k*. As shown in Figures 2g–h, a clear interaction emerged in the near-subthreshold regime: synchronization *R* exhibited a joint dependence on both *β* and *σ*, with higher *β* values requiring lower *σ* to achieve optimal enhancement. Notably, for each fixed *β, R* followed an inverted U-shaped function of *σ*, confirming the existence of an optimal noise intensity for every 1/f exponent. For example, optimal *σ* and local optimal *R* improvement pairs at representative values of *β* = (0.5,1.0,1.5,2.0) were: (1.7, 0.0453), (1.1, 0.0613), (0.6, 0.0702), (0.5, 0.0736), and (0.4, 0.0701) (Figure 2g, red circles). Moreover, when *σ* was fixed at its optimal value for each *β*, synchronization again exhibited an inverted U-shaped dependence on *β*, suggesting the existence of a global optimum resulting from the joint tuning of spectral structure and intensity.

To validate these findings, we extended the analysis to a 32-oscillator Kuramoto network and observed consistent results (Supplementary Materials II). Additional parameter sweeps, including variation in natural frequency ω, total iteration count *T*, and numerical step size, are also provided in the Supplementary Materials.

### tFNS Enhances Motor Cortex Synchronizations to Subthreshold Signals More Effectively than tRNS

To experimentally test whether exogenous noise with a biologically relevant 1/*f* structure facilitates subthreshold neural responses in the human brain, we conducted a study using the protocol illustrated in Figure 3a. In this paradigm, single-pulse transcranial magnetic stimulation (TMS) was delivered to the motor cortex. Suprathreshold TMS pulses elicit synchronous firing of corticospinal neurons at high frequencies^29^, producing detectable motor-evoked potentials (MEPs) in the contralateral target muscle. Consistent with our modeling framework, we used the probability of MEP occurrence—a macroscopic proxy for population-level neuronal synchrony— as a quantitative measure of cortical responsiveness.

**Figure 3.**
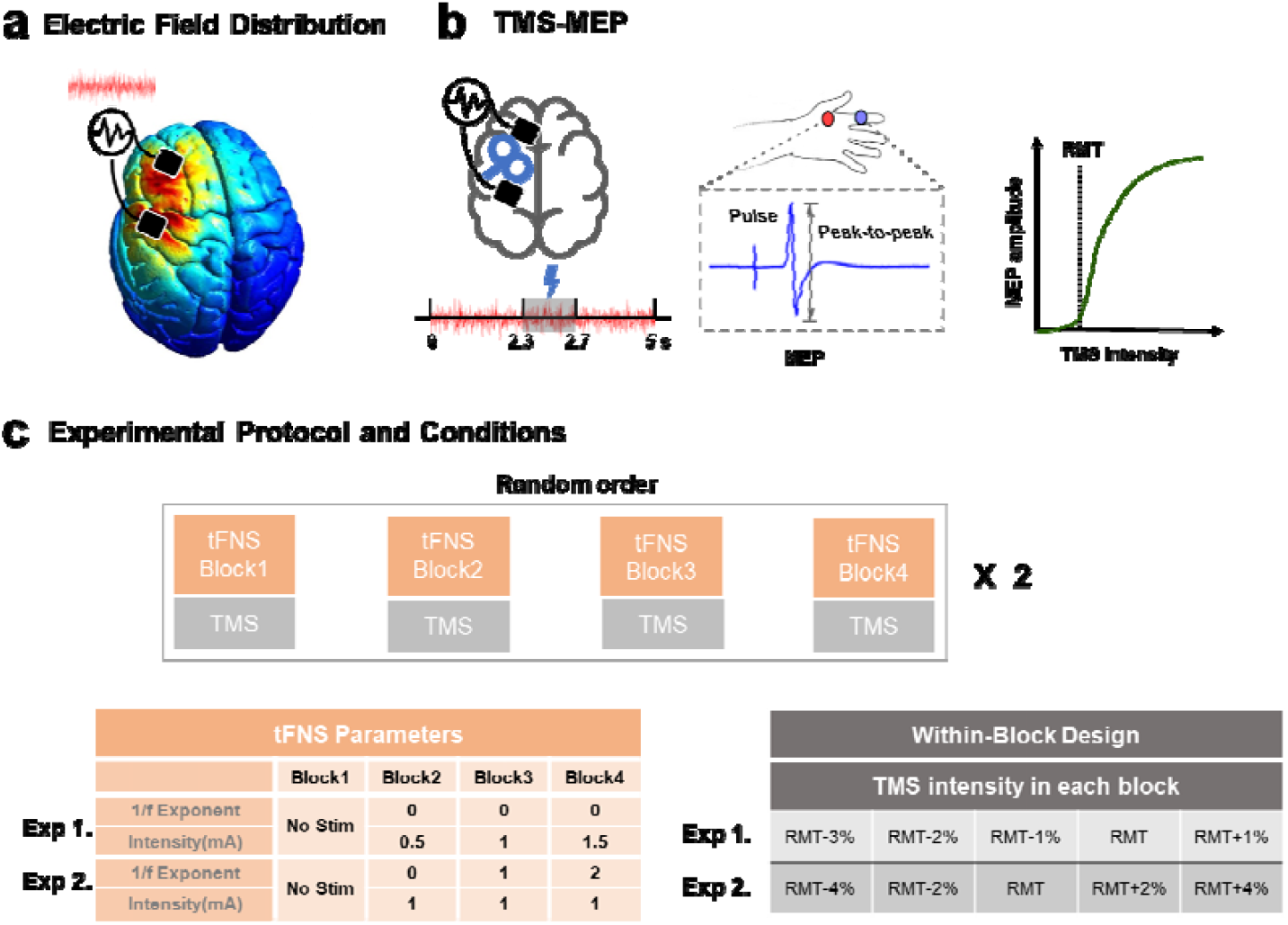
Experimental protocol and simulation of stimulation field in human brain experiments. **a**. Electric field simulations. Confirming cortical targeting, maximal fields localized in the area between stimulation electrodes F1 and CP3 (black rectangles). Color gradients reflect modeled electric field strengths. **b**. TMS-MEP diagram. TMS coil was placed in the “hotspot” for the first dorsal interosseous (FDI) muscle. MEPs were recorded from the right FDI muscle. Each trial lasted 5 s, with the TMS pulse randomly triggered between 2.3 and 2.7 s. The relationship between MEP amplitude and TMS intensities can be modelled using a sigmoid function. The RMT was defined as the minimum TMS intensity required to elicit MEPs of ≥50 □ μ V(peak-to-peak) of 10 trials. **c**. Stimulation protocol. Each experiment consisted eight stimulation blocks. TMS was delivered at five intensity levels in randomized order (10 trials per level) within each block. Randomly apply tRNS with varying parameters online in each block. In *Experiment 1*, tRNS with intensities of 0.5, 1.0, and 1.5 mA and 1/f exponent *β* = 0 (i.e., white noise) was applied across blocks, along with a no-stimulation control. TMS was delivered at five intensities: RMT– 3%, –2%, –1%, RMT, and RMT+1%. In *Experiment 2*, 1/*f* noise stimulation (tFNS) with *β* = 0, 1, 2 (common range in the human brain) at 1 mA was applied, along with a control condition. TMS intensities were RMT–4%, –2%, RMT, RMT+2%, and RMT+4%.

To probe the system’s sensitivity to subthreshold input, the TMS intensity was calibrated to just below the individual resting motor threshold (RMT), thereby ensuring that responses would be rare or absent without external modulation. We developed a transcranial 1/f noise stimulation (tFNS) system capable of delivering naturalistic electrical noise with adjustable intensity and spectral characteristics. We then applied it to the scalp region near the TMS site using two conditions: (1) variable noise intensity with fixed spectral profile (Figure 3b), and (2) variable 1/*f* spectral exponent at fixed intensity (Figure 3c), enabling simultaneous tFNS and TMS-MEP neuroimaging acquisition in humans. These transcranial 1/f noise stimulation (tFNS) protocols were compared to spectrally flat random noise stimulation (tRNS, *β* = 0) to evaluate their differential effects on cortical excitability.

### tRNS Enhances MEPs with an Optimal Intensity for Maximizing Cortical Synchronization

Motor-evoked potentials (MEPs) were recorded from peripheral muscles following TMS of the contralateral motor cortex. The resting motor threshold (RMT) was defined as the minimum TMS intensity required to elicit MEPs of ≥50 □ μV in at least 5 out of 10 trials. To examine the impact of transcranial random noise stimulation (tRNS) on corticospinal excitability, we measured the probability of evoking an MEP ≥50 □ μV (denoted as p(MEP_50μV_)) under five TMS intensity levels (RMT–3%, –2%, –1%, RMT, and RMT+1%) during 0.5, 1.0, and 1.5 □ mA tRNS, as well as a non-stimulation control.

A two-way mixed-effects model revealed significant main effects of both tRNS intensity (F_3,323_ = 8.81, p < 0.0001) and TMS intensity (F_4,323_ = 95.51, p < 0.0001), but no significant interaction (F_12,323_ = 1.19, p = 0.290). Post hoc comparisons with FDR correction showed that all tRNS intensities significantly increased p(MEP_50μ V_) relative to the control: 0.5 □ mA (p = 0.032, Cohen’s d = 0.43), 1.0 □ mA (p < 0.0001, d = 0.78), and 1.5 □ mA (p = 0.008, d = 0.51).

Focused analysis at RMT and its vicinity revealed that the facilitatory effects of tRNS were strongest near threshold (**Figure 4a**). At RMT–1% condition, a significant main effect of tRNS intensity was observed (F_3,51_ = 4.96, p = 0.004). Post hoc tests confirmed significant improvements in p(MEP_50μ V_) for all tRNS conditions compared to control: 0.5□mA (p = 0.012, d = 0.65), 1.0□mA (p = 0.010, d = 0.81), and 1.5 □ mA (p = 0.018, d = 0.64). Similarly, at RMT condition, a significant main effect of tRNS intensity was found (F_3,51_ = 4.94, p = 0.004), with post hoc results showing facilitation at 0.5 □ mA (p = 0.042, d = 0.66), 1.0 □ mA (p = 0.004, d = 0.98), and 1.5 □ mA (p = 0.042, d = 0.64). No significant enhancement was observed at intensities farther from threshold: RMT+1% (F_3,51_ = 2.17, p = 0.103), RMT–2% (F_3,51_ = 1.05, p = 0.378), or RMT–3% (F_3,51_ = 0.96, p = 0.418).

**Figure 4.**
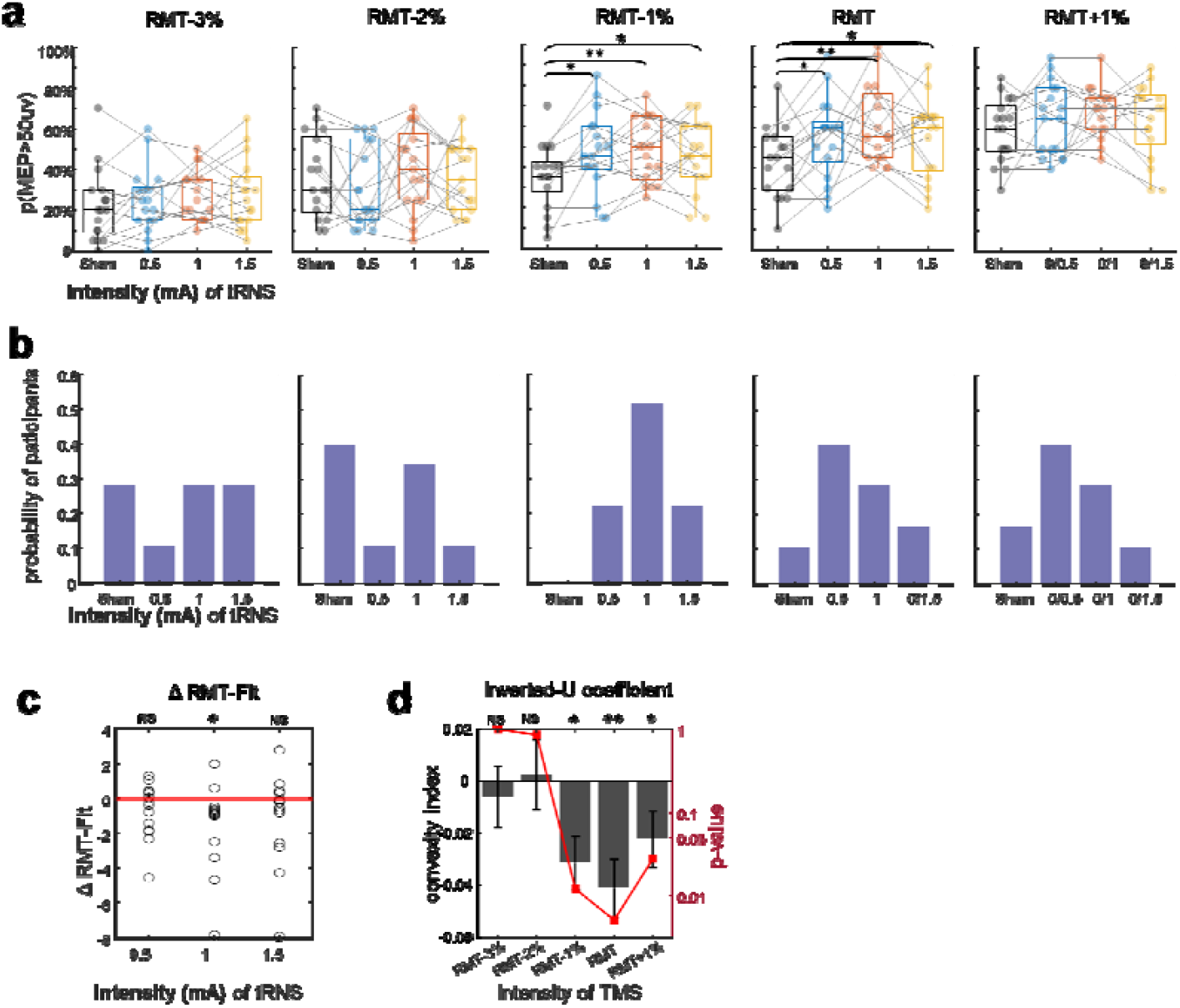
Enhancement of MEP probability by tRNS. **a**. Probability of evoking MEPs ≥50μV p(MEP_50μ V_) across TMS intensities (RMT–3% to RMT+1%) under no stimulation (control) and tRNS at 0.5, 1.0, and 1.5 mA. Individual data points and group means are shown. Two-way mixed-effects model; FDR-corrected. **b**. Distribution of optimal tRNS intensity per participant across TMS levels, defined as the tRNS intensity yielding maximal p(MEP_50μ V_). **c**. RMT-fit values for each condition. Mixed-effects model; FDR-corrected. **d**. Convexity index (CI) values across TMS intensities. CI < 0 indicates an inverted-U relationship between tRNS intensity and p(MEP_50μ V_). Significance tested against zero distribution. Bars represent mean ± SEM.

To further quantify tRNS effects, we refitted the threshold (RMT-fit value) for each tRNS intensity. A main effect of tRNS intensity on RMT-fit was observed (F_3,51_ = 3.17, p = 0.034), with post hoc analysis revealing a significant reduction only for 1.0 □ mA tRNS (p = 0.039, d = 0.64; **Figure 4c**). Among all intensities, 1.0 □ mA tRNS produced the strongest enhancements at both RMT (p = 0.004, d = 0.98) and RMT–1% (p = 0.010, d = 0.82) condition, suggesting a well-defined optimal stimulation level.

Given the high inter-individual variability in MEP responses, we examined the optimal stimulation intensity at the individual level (**Figure 4b**). For both RMT and RMT–1% conditions, most participants showed maximal benefit at 0.5 or 1.0 □ mA tRNS. In contrast, at RMT–2% condition, the largest proportion of participants responded best to the no tRNS control, indicating that adding noise to deeply subthreshold input can reduce detectability. At RMT+1% condition, the distribution of optimal intensities resembled that at RMT, suggesting that moderate facilitation may extend marginally into the suprathreshold regime.

Finally, we applied the convexity index to assess whether the relationship between p(MEP_50μ V_) and tRNS intensity followed an inverted-U shape. As shown in **Figure 4e**, CI values were significantly less than zero at RMT–1% (p = 0.012), RMT (p = 0.005), and RMT+1% (p = 0.028) conditions, indicating a robust convex relationship. These results confirm the existence of an **optimal tRNS intensity** that can reliably enhance neural synchrony to near-threshold TMS input.

### tFNS Outperforms tRNS in Enhancing Corticospinal Excitability and Expands Effective Range

To test whether scale-free noise more effectively facilitates corticospinal excitability, we delivered tFNS with 1/f exponents *β* = 0, 1, and 2 at 1 mA, and measured p(MEP_50μ V_) across five TMS intensities (RMT–4% to RMT+4%). A two-way mixed-effects model revealed significant main effects of tFNS condition (F_3,294_ = 7.20, p < 0.0001) and TMS intensity (F_4,294_ = 158.42, p < 0.0001), with no significant interaction (F_12,294_ = 0.63, p = 0.818). Post hoc tests (FDR-corrected) indicated that *β* = 2 significantly increased p(MEP50_μ_V) versus both control (p < 0.0001, d = 0.66) and *β* = 0 (p = 0.091, d = 0.38). *β* = 1 also significantly improved outcomes over control (p = 0.032, d = 0.58).

At RMT condition, tFNS significantly boosted p(MEP50_μ_V) (F_3,46_ = 5.35, p = 0.003) and reduced RMT-fit (F_3,46_ = 9.03, p < 0.0001) (**Figure 5a, c**). Post hoc analysis revealed tFNS with *β* = 0 boosting p(MEP_50μ V_) (p = 0.016, Cohen’s d = 0.61), and reduced RMT-fit (p = 0.021, Cohen’s d = 0.66) compared to control. As tFNS with *β* = 0 is tRNS, these results verified the effect of 1mA tRNS in Experiment 1. Notably, tFNS with both *β* = 1 and 2 produced larger effect sizes than *β* = 0 at RMT condition (*β* = 1, p=0.011, d = 0.55; *β* = 2, p=0.007, d = 0.93) and reducing RMT-fit (*β* = 1: p < 0.0001, d = 0.9, *β* = 2: p < 0.0001, d = 0.82).

**Figure 5.**
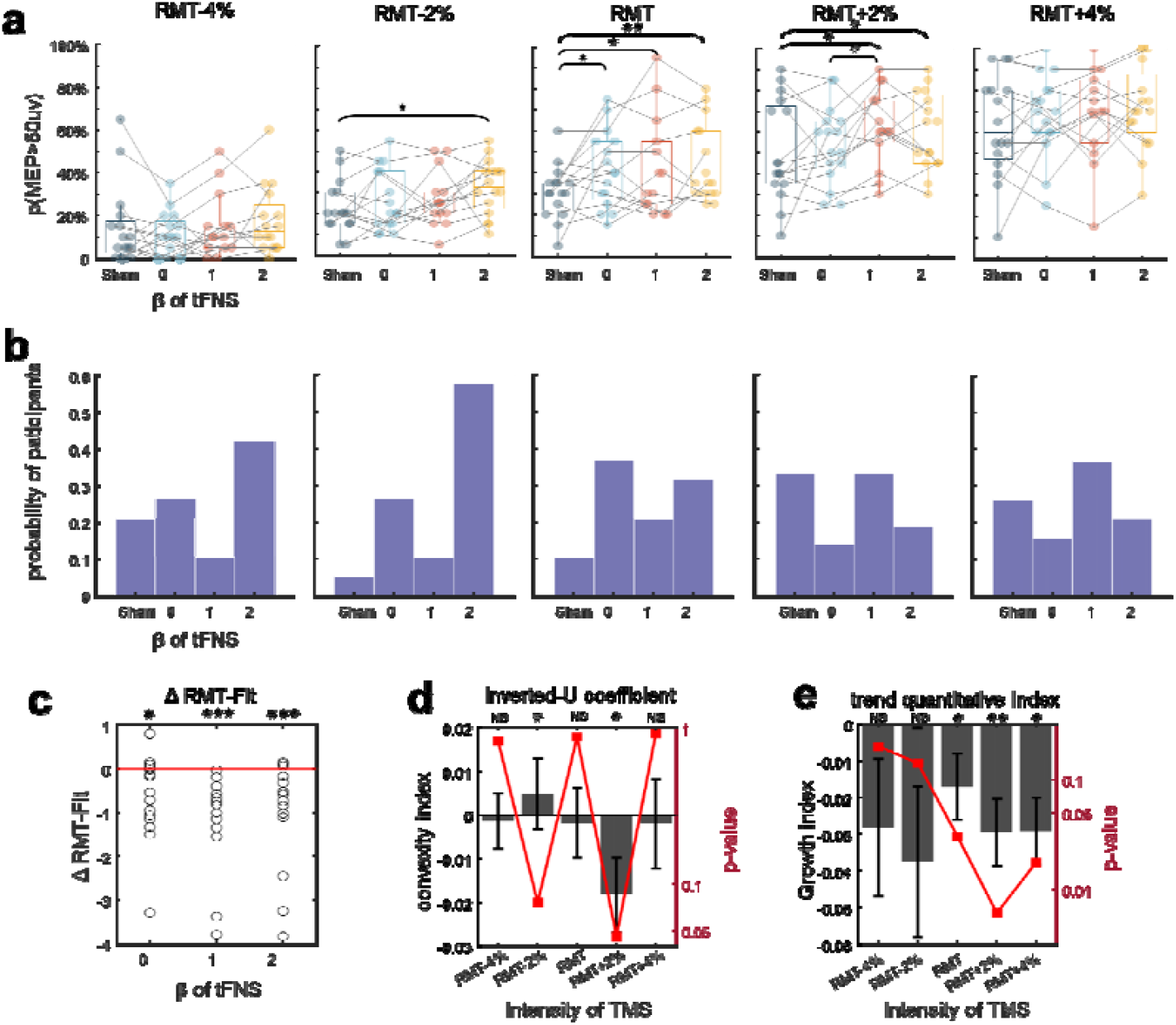
Enhancement of MEP probability by tFNS across 1/f exponents. **a**. p(MEP_50μ V_) under tFNS with 1/f exponents *β* = 0 (tRNS), 1, and 2, at 1 mA intensity, compared to no-stimulation control. TMS intensities: RMT–4% to RMT+4%. **b**. Distribution of individual optimal *β* values per participant across TMS levels, defined as the *β* yielding maximal p(MEP50μV). **c**. RMT-fit across *β* conditions. Mixed-effects model; FDR-corrected. **d**. Convexity index values across TMS intensities. CI < 0 indicates inverted-U trend in *β* vs. p(MEP_50μ V_); significance tested against zero. **e**. Growth index analysis of p(MEP_50μ V_) across increasing *β*, where positive values indicate upward trends. Bars represent mean ± SEM.

**Figure 6.**
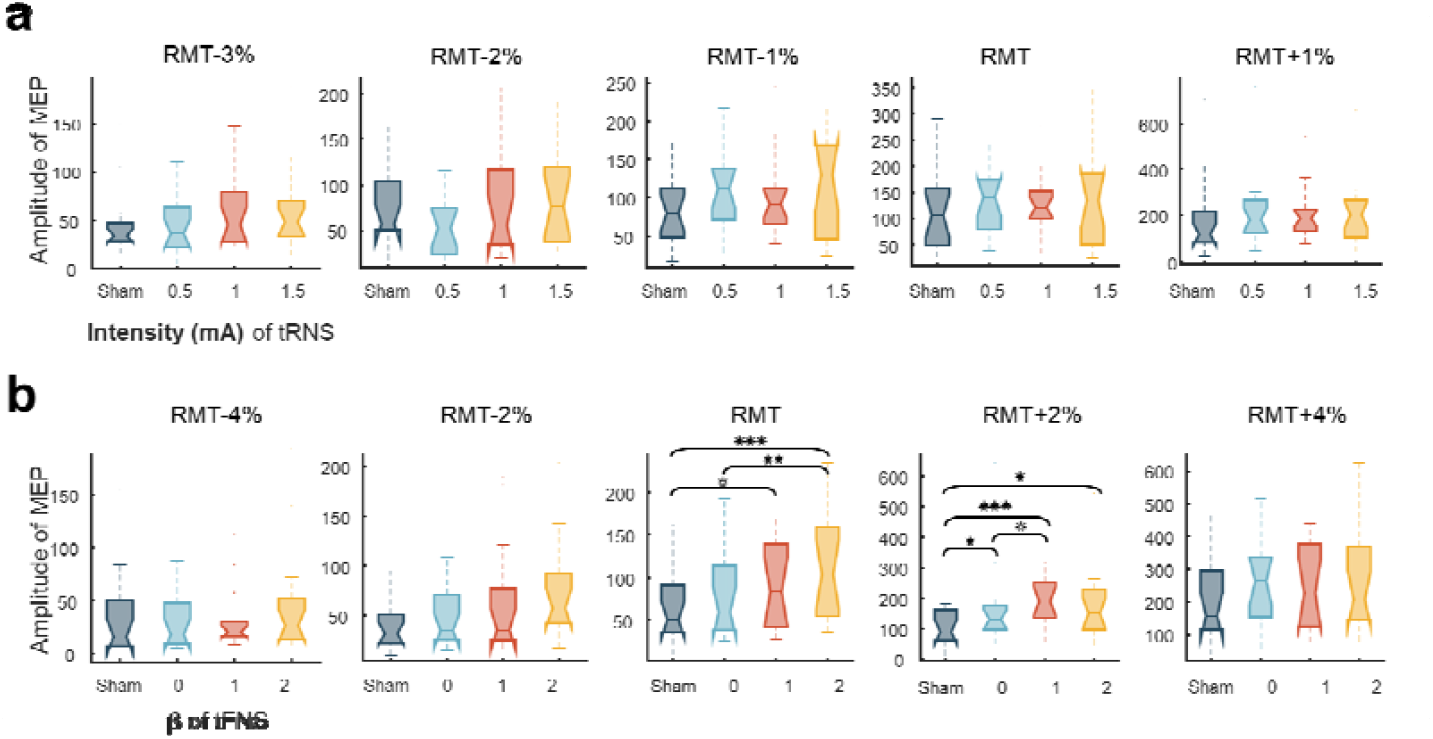
MEP amplitude changes under tRNS and tFNS. **a**. Mean and individual MEP amplitudes in Experiment 1 (tRNS). No significant group-level effect. **b**. MEP amplitude enhancement in Experiment 2 (tFNS). *β* = 1 and 2 significantly improved amplitude over *β* = 0 and no stimulation. Mixed-effects model; FDR-corrected.

Beyond threshold, tFNS also enhanced p(MEP_50μ V_) at RMT–2% (F_3,46_ = 3.52, p = 0.022) and RMT+2% (F_3,46_ = 4.14, p = 0.011). These effects were driven by *β* = 1 (RMT+2%: p = 0.021, d = 0.61) and *β* = 2 (RMT-2%: p = 0.020, d = 0.91). Error! Reference source not found.**b** showed the distribution of optimal 1/f exponent for each participant. The optimal 1/f exponents are concentrated on *β* = 2 at RMT-4% and RMT-2%condition. For subthreshold TMS, the optimal parameters are concentrated on *β* = 1 and control.

Importantly, convexity index analysis showed significant inverted-U patterns at RMT+2% (p = 0.046; **Figure 5d**), suggesting optimal spectral structure in the suprathreshold regime. This is consistent with the inverted U-shaped pattern observed on the weakly suprathreshold in the Kuramoto model 2, further demonstrating that 1/f noise has less destructive effect on the weakly suprathreshold stimulation. Growth index analysis (**Figure 5e**) revealed a monotonic increase in MEP probability with increasing *β* at RMT (p = 0.03), RMT+2% (p = 0.006), and RMT+4% (p = 0.018), indicating *β* = 2 may not yet represent the peak enhancement point.

Critically, tFNS also enhanced MEP amplitude, a proxy of corticospinal output strength. In **Experiment 1**, a two-way linear mixed-effects model was used to assess the impact of tRNS (white noise) and TMS intensity on the amplitude of evoking motor responses. The analysis revealed a significant main effect of TMS intensity (F_4,323_ = 62.80, p < 0.0001), but neither a significant main effect of tRNS intensity (F_3,323_ = 2.10, p = 0.10) nor an interaction effect (F_12,323_ = 0.74, p = 0.818). These findings are consistent with previous reports showing limited group-level effects of white-noise stimulation on corticospinal excitability^26^.

By contrast, Experiment 2 demonstrated that transcranial 1/f noise stimulation (tFNS) significantly modulates both MEP probability and amplitude. The two-way model showed robust main effects of both tFNS condition (F_3,263_ = 12.93, p < 0.0001) and TMS intensity (F_4,29_ = 14.18, p < 0.0001), along with a significant interaction (F_12,263_ = 1.96, p = 0.028). Post hoc comparisons (FDR-corrected) confirmed that tFNS with all tested exponents significantly increased the amplitude of evoking an MEP compared to the no-stimulation control: *β* = 0: p = 0.015, Cohen’s d = 0.49; *β* = 1: p < 0.0001, d = 0.85; *β* = 2: p < 0.0001, d = 0.86.

At the RMT level, a significant main effect of *β* was observed (F_3,46_ = 7.45, p < 0.001), with *β*□= □2 producing significantly greater responses than both *β* □ =□0 (p = 0.007, d = 0.68) and the no-stimulation condition (p < 0.001, d = 1.28). At RMT+2%, all tFNS conditions led to significantly increased amplitudes relative to baseline: *β* = 0: p = 0.036, d = 0.54; *β* = 1: p = 0.0006, d = 0.87; *β* = 2: p = 0.015, d = 0.72. The pairwise comparison between *β*□= □1 and *β*□= □0 approached significance (p = 0.094, d = 0.36), suggesting a graded enhancement trend with increasing 1/f exponent.

Together, these results establish that scale-free noise stimulation—particularly with higher 1/f exponents—more effectively enhances corticospinal excitability than spectrally flat noise. This supports the hypothesis that the functional engagement of neural circuits is improved when exogenous noise matches the endogenous aperiodic structure of brain activity^3, 30, 31, 32, 33, 34^.

## Discussion

In this study, we employed a confirmatory modeling and experimental approach to investigate how noise with biologically inspired spectral structure modulates neural synchrony. Specifically, we introduced a scale-free exponent (*β*) as a defining parameter of 1/*f* noise and validated its impact through both numerical simulations and human TMS-MEP experiments. Our results demonstrate that transcranial random noise stimulation (tRNS) at an optimal intensity enhances the detection of near-threshold TMS stimuli, and that this effect is further amplified when the stimulation adopts an endogenous-like 1/*f* spectral structure (tFNS). The enhancement extended to both the probability and amplitude of motor evoked potentials (MEPs), and across a broader range of input intensities. Consistent with our Kuramoto model simulations, we observed that optimal neural enhancement arises from a **competitive synergy** between noise intensity and 1/*f* exponent. These findings provide mechanistic insight into how scale-free stimulation can entrain and augment neural dynamics associated with the brain’s aperiodic components.

### Synergistic Effects Between Noise Intensity and 1/f Structure

In Experiment 1, we confirmed that white-noise tRNS reduces motor cortex threshold, in line with earlier findings where tRNS facilitated detection of subthreshold sensory inputs^20^ or reduced RMT with 2□mA stimulation^26^. The presence of an optimal noise intensity was validated using a convexity index, supporting the framework of stochastic resonance in neural systems^20, 22, 35^. Our simulations provide a mechanistic basis: in weakly coupled oscillator systems, noise exhibits an inverted U-shaped relationship with synchronization^3^, while in overly coupled systems, added noise disrupts synchrony. This implies that excessive noise in an already-optimized system can degrade output—an idea consistent with traditional views of noise as detrimental. However, prior studies failed to observe optimal tRNS intensity effects^26^, possibly due to coarse sampling or inter-individual variability in thresholds. In contrast, our finer sampling and modeling revealed a reliable optimal regime.

Importantly, our study first demonstrated that 1/f noise outperforms white noise in several ways: stronger enhancement of MEP probability, broader effective TMS intensity ranges, increased MEP amplitude, but also imposes less destructive interference in suprathreshold signals. These findings support our hypothesis that stimulation with naturalistic statistical structure more effectively engages the brain’s intrinsic dynamics. This aligns with previous work showing that neurons preferentially respond to input statistics that mirror endogenous fluctuations ^30, 33, 36, 37^—such as the auditory system’s sensitivity to 1/*f* structure in natural sounds^37^, or improved memory for sequences with scale-free temporal properties^33^ and 1/f noise preserve the quantum criticality^34^.

### Antagonism and Optimality Between Noise Parameters

Unexpectedly, an inverted U-shaped dependency on the 1/*f* exponent (*β*) was observed only for suprathreshold TMS stimuli. This asymmetry may be explained by two factors: (1) the limited resolution in our sampling of *β* values, or (2) the need for parameter coordination between intensity and spectral structure. Our model simulations supported the latter. They reveal a synergistic antagonism—increased *β* enhances synchronization when paired with reduced intensity, culminating in a global optimum. This interaction suggests that optimizing neural responses requires not just tuning a single noise parameter but balancing both intensity and spectral slope. Previous studies have also highlighted competitive or synergistic effects of multiple system parameters, such as the optimal balance between loss and noise inducing bistability in nonlinear resonator systems ^38^, and the balance between noise and coupling connectivity controlling cell differentiation^39^. Such fine-tuned interactions may generalize to other sensory and cognitive domains involving weak signal detection, such as perceptual awareness or attention.

### Potential Mechanisms: Resonance With Intrinsic Neural Dynamics

The superior efficacy of 1/*f* noise likely stems from its resonance with the brain’s endogenous dynamics^16^. Neural activity exhibits a 1/*f* power spectrum, thought to reflect multi-scale coordination and excitation–inhibition (E/I) balance^40^. Steeper spectral slopes have been linked to inhibitory (GABAergic) tone, while flatter spectra reflect higher excitatory activity [37]. In disorders of consciousness, aging, and sleep, a flatter 1/*f* slope is associated with E/I imbalance^41^. We speculate that 1/*f* stimulation may entrain and restore intrinsic aperiodic structure, rebalancing E/I activity and improving responsiveness to subthreshold stimuli^40^.

Additionally, 1/*f* dynamics are a hallmark of self-organized criticality. The brain operating near criticality avoids both hyper-excitation and quiescence, enabling optimal information processing^42^. Our findings suggest that stochastic perturbation with scale-free properties may push neural systems toward such critical states, resulting in larger and more stable macroscopic responses^11, 34^. Prior modeling research has shown 1/*f* noise facilitates quantum critical transitions^34, 43^. Previous experimental studies also have reported noise-induced ordered macroscopic responses^13, 44^ : long tail distribution of neural avalanches^45, 46^, traversals of dynamic repertoire^13^ and bistability of gene expression^44, 47^ and stable cellular differentiation^39^. In line with this, noise-induced enhancements in our model occurred near phase transition thresholds.

Thus, while brain activity is influenced by microscale randomness, our data indicate that global order can emerge from parameterized stochastic modulation—in this case, through the structured application of 1/*f* noise to suppress noise levels, maintaining the brain on the edge between order and disorder for efficient information processing.

### Clinical Implications: A Novel Path for Non-Invasive Neuromodulation

Our findings open new avenues for clinical translation of 1/*f*-based neuromodulation. While the 1/*f* structure has been implicated in neurological and psychiatric disorders^15^, few studies have attempted to entrain or restore this structure via external stimulation. A recent fMRI study demonstrated that transcranial alternating current stimulation (tACS) could restore avalanche-like 1/*f* dynamics in Parkinson’s disease, improving behavioral outcomes.

tFNS may represent a next-generation neuromodulation approach capable of restoring critical-like dynamics in patients with impaired consciousness or age-related cognitive decline. By tuning the 1/*f* exponent to better match endogenous fluctuations, tFNS could help reestablish optimal operating conditions in disordered neural networks.

## Conclusion

Simulations using the Kuramoto model revealed a global optimal enhancement of synchronization when increasing 1/*f* exponent while decreasing noise intensity. This antagonistic interaction suggests that optimal system performance is achieved by carefully balancing these parameters. Human TMS-MEP experiments confirmed this synergy, showing that 1/*f*-structured noise (tFNS) amplified both the probability and amplitude of evoked motor responses beyond what was achievable with white-noise tRNS alone. These findings offer new insight into the constructive role of biologically structured noise in brain function. By showing how 1/*f* properties can be leveraged to suppress internal noise while enhancing coordination, we propose a principled strategy for maintaining neural systems on the edge between order and disorder—an optimal state for adaptive information processing.

## Materials and Methods

### Kuramoto Model Simulations

The Kuramoto model was used to simulate phase synchronization among coupled neural oscillators. This model describes the evolution of oscillator phases in systems characterized by nonlinear coupling and has been widely applied in neuroscience to study network synchronization^4^.

Model1: We first analyzed a two-oscillator system under Gaussian white noise of varying intensity:

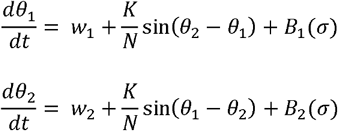

where θ(t) are oscillator phases, ω are the natural frequencies, *K* is the coupling constant, N is the number of oscillators, and B are independent Gaussian white noise inputs (*U*(0, *σ* ^2^))). We fixed w_1_=2.5, w_2_=4.5, and explored effective coupling by normalizing *k*=*K*/(w_2_−w_1_), ranging from 0 to 2, step by 0.05 (*K*_∈_[0,4], step by 0.1), with simulations focusing on *k*≤1.5. The synchronization index *R* was computed from the complex order parameter: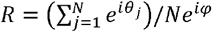, *R* was averaged across 1500 simulation steps.

Model2: To examine the effects of 1/f noise, we replaced the white noise term with filtered noise of a defined 1/f spectral exponent *β*:

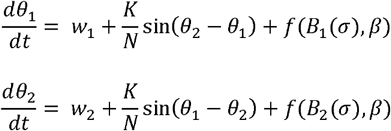

where, *f*(_⋅_) denotes spectral filtering to generate 1/*f* noise. We set w_1_=2 and w_2_=5 with *k*=*K*/(ω2−ω1)_∈_[0,4/3] (*K*_∈_[0,4], step by 0.1). Simulations were performed for fixed intensity *σ*=1 while varying 1/f exponent *β*_∈_[0,2] in steps of 0.1. We then tested interaction effects by varying both *σ*_∈_[0,4] and *β*_∈_[0,2].

Full derivations and filtering implementation are described in **Supplementary Materials II**.

### Transcranial Noise Electrical Stimulation (tRNS and tFNS)

A custom-built stimulation device delivered white (tRNS) or 1/f noise (tFNS) through 20 mm Ag/AgCl electrodes with conductive gel (GT10, Greentek, China). Electrode positions followed the 10-10 system, secured by elastic caps. The stimulation current ranged from 0.5 to 1.5 mA (peak-to-baseline), corresponding to a current density of 0.16–0.48 mA/cm^2^. System designs are provided in **Supplementary Materials I**.

### Electric Field Modeling

Finite element simulations were performed using SimNIBS 4.0^48^, based on the MNI152 template^49^. Electrodes (2 cm diameter, 3 mm thickness) were placed at F1 and CP3 according to the 10-10 electrode system. A current of 1 mA was applied. The electric field was concentrated within the cortical region between F1 and CP3, confirming delivery to motor-related areas.

### Transcranial Magnetic Stimulation (TMS)

Transcranial Magnetic Stimulation (TMS), introduced by Barker et al. in 1985, is a non-invasive neurophysiological tool used to investigate functional characteristics of the human motor cortex-spinal cord pathway. The principle of TMS involves the generation of a transiently changing magnetic field by applying a brief, time-varying current pulse to a stimulating coil. According to Faraday’s law of electromagnetic induction, this changing magnetic field induces an electric current in cortical neurons. According to Faraday’s law of electromagnetic induction, this changing magnetic field induces an electric current in cortical neurons.

In this study, TMS was delivered using a Magstim Rapid stimulator with a 70 mm figure-of-eight coil. The coil was positioned tangentially to the scalp surface, 45° to the midline, with the handle pointing backward, targeting the “hotspot” for the first dorsal interosseous (FDI) muscle. This site was determined as the point evoking the most reliable MEPs and marked for reliable consistent placement.

### Electromyography (EMG) Signal Recordings and Processing

The cortical-spinal neurons could be depolarized by TMS at the motor cortex, leading to muscle controlled by stimulated cortical region activating. These TMS-evoked muscle activations, known as Motor Evoked Potentials (MEPs), can be recorded using electromyography (EMG). EMG was recorded from the right FDI muscle using a custom amplifier (bandwidth: 0.16–658 Hz; sampling rate: 2000 Hz). TMS pulses were synchronized via a trigger signal.

The MEP amplitude (peak-to-peak) was extracted using a custom MATLAB script. Resting motor threshold (RMT) was defined as the TMS intensity eliciting MEPs ≥50 µV in ≥50% of trials. To estimate RMT-fit^26^ for each tRNS/tFNS condition, a linear fit (y = ax + b) was applied to the relationship between TMS intensity (x) and p(MEP_50μ V_) (y). The threshold was computed as RMT-fit=(0.5−b)/a. Participants whose data could not be reliably fit (3 in Exp. 1, 1 in Exp. 2) were excluded.

### Experimental Design

Experiment 1 – tRNS. In the first experiment, we tested the online effect of white noise electrical stimulation on corticospinal excitability. Twenty-four right-handed participants were recruited. Before the formal experiment, we conducted a preliminary test to determine the resting movement threshold (RMT) of each participant. After RMT calibration, seven were excluded due to unstable thresholds, leaving 17 participants (9 females, age = 23□±□2.23 years). Each participant underwent 8 blocks: TMS intensities (RMT±1–3%) were presented in randomized order (10 trials/intensity), and tRNS was applied at 0, 0.5, 1.0, or 1.5□mA in randomized blocks (2 blocks/condition, 20 trials/condition per TMS level). Each trial took 5s as the window, and the TMS stimulus was randomly triggered within 2.3-2.7s from the start of the trial.

Experiment 2 – tFNS. Nineteen right-handed participants were tested using 1 mA tFNS with *β* = 0, 1, 2, and no tFNS control condition. None of the participants participated in both of experiment 1 and 2. After excluding three participants due to threshold inconsistencies, 16 were included (9 females, age = 23.69□±□2.44 years). TMS intensities were RMT±2–4%. Two participants did not receive *β* = 1 due to discomfort. Other procedures were as in Experiment 1.

### Trend Quantification Analysis

To quantify non-monotonic and growth trends in stimulation effects, we developed two metrics:

**Convexity Index (CI)**: Captures inverted-U structure in discrete data sequences:

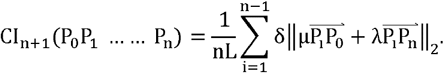

CI □< □0 indicates an inverted-U trend; higher magnitude reflects stronger curvature. Properties and derivation are in **Supplementary Materials III**.

### Growth Index (GI)

Measures monotonic increase. GI □> □0 indicates increasing trend; larger values indicate steeper growth. The combination of these two indices can effectively analyze the trends of various discrete data.

### Statistical Analysis

Linear Mixed Effects Models (LME) were used to analyze MEP probability and RMT-fit, implemented using the lme4 package in R (v4.3.1). Model structure:

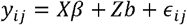

where: y_ij_: response for subject i at trial j; *β*: fixed effects (e.g., stimulation, TMS intensity); b: random effects (e.g., subject variability); □_ij_: residual error.

In Experiment 1, fixed effects: tRNS intensity (sham, 0.5, 1.0, 1.5 mA), TMS intensity (RMT±1– 3%) and interaction effect of them. In Experiment 2, fixed effects: *β* = 0, 1, 2 (plus sham), TMS intensity (RMT±2–4%) and interaction effect of them. ANOVA was used for fixed effects. Models were estimated via restricted maximum likelihood (REML), and model fit was assessed via Akaike Information Criterion (AIC). Post hoc comparisons were FDR-corrected. Cohen’s d was used to estimate effect sizes. In addition, we conducted a confirmatory analysis of the simple effects of tRNS for each TMS intensity using LME. Both convexity and growth indices were tested against 0 using one-sample t-tests.

## Author Contributions

Anan Ping: conceptualization, methodology, investigation, visualization, formal analysis, writing original draft, writing—review and editing, validation, data curation; Longzhou Guan: methodology, software, validation; Hai Gu: methodology, investigation, formal analysis, visualization, writing—review and editing; Zhipeng Jiang: software, validation; Wenhao Deng: methodology, software,; He Chen: conceptualization; Ke Zeng: writing—review and editing; Xiaoli Li: conceptualization, investigation, writing original draft, writing—review and editing, validation.

## Funding

This work was supported by STI2030 Major Projects+2021ZD0204300 (to X. Li).

## Conflicts of Interest

Authors declare that they have no competing interests.

## Data and code availability

All data, codes, and materials used in the analysis will be available in the future.

## Notes

### Competing Interest Statement

The authors have declared no competing interest.

