## Supplementary Materials for "Enhancing Neural Synchrony with Endogenous-like 1/f Noise Stimulation"

### Ⅰ. 1/f Noise Generation and Output

#### A. Generation of 1/f noise with different power-law exponent

1/f noise (also known as power-law noise) is a type of colored noise that is ubiquitous in many natural and man-made systems. The characteristic of power-law noise is that its power spectral density is proportional to the reciprocal of frequency, i.e.,$S(f)\propto1/f^{\beta}$, where β is the inverse power of frequency. Typically, in power spectrum representations, 1/f signals are parameterized by an offset and an exponent. For a specific exponent, the offset is determined by the signal intensity.

In this study, power-law noise signal was generated by filtering an initial Gaussian white noise (GWN) signal. The intensity of this base GWN signal was controlled via a set peak current parameter. This parameter is statistically defined as 1.96 times the signal's standard deviation (σ), ensuring that the instantaneous output current remains below this peak value with 95% probability. To introduce the specific power-law spectral characteristics, this GWN signal was processed through a colored filter. Following the methodology described in literature^[1]^, we implemented this filtering stage using a 63rd-order autoregressive (AR) model. The coefficients for this AR model were determined by the following formula:

$$a_{0}=1，a_{k}=(k-1-\frac{\beta}{2})\frac{a_{k}-1}{k}, k=1,2,\ldots,63.$$

For special noise generation with β=1 (pink noise) and β=2 (Brown noise), this is achieved by specially tuned second-order knot (SOS) filters, where pink noise is generated by 6 second-order filters (total order 12) and Brown noise is generated by 5 second-order filters (total order 10), which are optimized to provide better performance. The above process can be implemented by the dsp.ColoredNoise function of MATLAB.

The 1/f noise signal exhibits increasing energy with decreasing frequency. In practical physical systems, to prevent output current saturation and confine the signal bandwidth within the effective system bandwidth, a 5Hz-700Hz bandpass filter (12th-order, Chebyshev Type-I, IIR filter) is employed.

Since 1/f noise signals are generated by filtering Gaussian white noise, their Power Spectral Density (PSD) distribution is altered, consequently affecting the signal's total power. Therefore, when configured using the same peak current parameter, 1/f noise with different spectral exponents (β) and tRNS (Gaussian white noise, β=0) will deliver different levels of actual power. This results in a discrepancy in the total energy injected over a fixed time period for stimuli with identical peak current settings but different β values.

To ensure equivalent energy delivery across different noise types (i.e., noise with varying β values) under identical peak current settings, we implemented a power normalization procedure. This approach uses the power of the standard tRNS signal (Gaussian white noise) as a reference. For a 1/f noise signal with a specific β value, its amplitude is adjusted by multiplying with a pre-calculated gain factor (Gain Factor) to match its total power to that of the tRNS signal configured with the same peak current.

This gain factor was determined empirically using MATLAB. For each specific β value, a 1/f noise signal was generated, and its Root Mean Square (RMS) value (representing the effective signal value) was calculated over a predefined time window (e.g., 25 seconds). This RMS value was then compared to the RMS value of a standard Gaussian white noise signal generated subject to the same peak current constraint (1.96σ). The gain factor was computed as the ratio of the RMS of the white noise to the RMS of the colored noise (Gain = RMS_white_ / RMS_colored_), thereby compensating for power variations introduced by spectral shaping.

#### B. Design of transcranial random electrical noise equipment

The electrical stimulation device employs the ESP32-PICO-D4 (Espressif Systems (Shanghai) Co., Ltd., China) chip as the main controller, and a 16-bit DAC (DAC8562, Analog Devices Inc., USA) for high-precision signal output. A high-voltage operational amplifier (OPA454, Texas Instruments Inc., USA) forms the voltage-controlled current source circuit, generating the stimulation current. The device is capable of delivering high-precision current at a sampling rate of 10 kSps, with a current precision of ±3 μA, a maximum output voltage of ±25 V, and an analog bandwidth ranging from DC to 1.6 kHz.

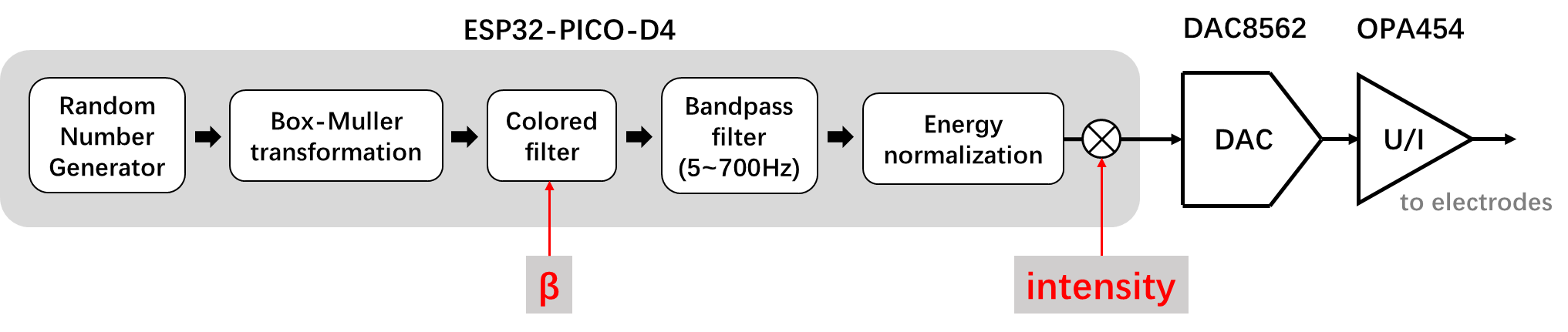

Figure S1 System Architecture and Signal Generation

The MCU's random number generator can produce uniformly distributed random numbers. By applying the Box-Muller transformation to this signal, Gaussian white noise with a normal distribution in the time domain and a uniform distribution in the frequency spectrum can be obtained. Gaussian white noise is used to generate 1/f noise as described in Section A of Part Ⅰ.

### Ⅱ. Model simulation and analysis

#### A. Iterative approach

For different oscillators $\theta_{i}$, when noise $B_{i}$ is respectively applied to each，its Kuramoto model can be described as follows：

$\frac{d\theta_{i}}{dt}= w_{i}+\frac{K}{N}\sum_{j=1}^{N} \sin\left( \theta_{j}-\theta_{i} \right)+B_{i} .$	(S1)

The complex-valued order parameter is defined as $Re^{i\varphi}= (\sum_{j=1}^{N} e^{i\theta_{j}})/N$. Multiply both sides of the formula by $e^{-i\theta_{i}}$ to get：$Re^{i(\varphi-\theta_{i})}= (\sum_{j=1}^{N} e^{i{(\theta}_{j}-\theta_{i})})/N$. Expand it using Euler's formula, pay attention to its imaginary part: $Rsin(\varphi-\theta_{i})=\sin\left( \theta_{j}-\theta_{i} \right)/N$, substitute this formula into Kuramoto model to obtain:

$\frac{d\theta_{i}}{dt}= w_{i}+KRsin(\varphi-\theta_{i})+B_{i}.$	(S2)

Further, in order to express R explicitly with $\theta$, modulo both sides of the complex-valued order parameter, we can get:

$R=\sqrt{\left( \frac{\sum_{j=1}^{N} cos\theta_{j}}{N} \right)^{2}+\left( \frac{\sum_{j=1}^{N} sin\theta_{j}}{N} \right)^{2}}.$	(S3)

Meanwhile, consider the real and imaginary parts of the complex-valued order parameter $Rcos(\varphi)=(cos\theta_{1}+cos\theta_{2})/2，Rsin(\varphi)=(sin\theta_{1}+sin\theta_{2})/2$, and use $Rsin(\varphi)$ divided by $R+Rcos(\varphi)$ and simplify to explicitly express $\varphi$ in terms of $\theta$:

$\varphi=2arctan\left( \frac{\sum_{j=1}^{N} sin\theta_{j}}{NR+\sum_{j=1}^{N} cos\theta_{j}} \right).$	(S4)

Thus, combining the aforementioned equations and using the Euler method for numerical solution^[2, 3]^, an iterative formula can be obtained^[1]^：

$\theta_{i}(n)=\theta_{i}(n-1)+K*R(n-1)*sin(\varphi(n-1)-\theta_{i}(n-1))*dt+w_{i}*dt+B_{i}*\sqrt{dt} , i=1,2,\ldots,N$. (S5)

where, $K \mathrm{and}w_{i}$ are parameters, taken as:

$$dt=0.05, R(n-1)=\sqrt{\left( \frac{\sum_{j=1}^{N} sin\theta_{j}(n-1)}{N} \right)^{2}+\left( \frac{\sum_{j=1}^{N} sin\theta_{j}(n-1)}{N} \right)^{2}}, \varphi(n-1)=2arctan\left( \frac{\sum_{j=1}^{N} sin\theta_{j}(n-1)}{NR+\sum_{j=1}^{N} cos\theta_{j}(n-1)} \right).$$

Considering the case of a coupled-oscillator model (N=2), the iterative formula can be simplified to:

$\left\{ \begin{aligned} \theta_{1}(n)=\theta_{1}(n-1)+K*R(n-1)*sin(\varphi(n-1)-\theta_{1}(n-1))*dt+w_{1}*dt+B_{1}*\sqrt{dt} \\ \theta_{2}(n)=\theta_{2}(n-1)+K*R(n-1)*sin(\varphi(n-1)-\theta_{2}(n-1))*dt+w_{2}*dt+B_{2}*\sqrt{dt} \end{aligned} \right.$	(S6)

where, $K,w_{1},w_{2}$ are parameters, taken as:

$$dt=0.05, R(n-1)=\sqrt{\frac{1+cos(\theta_{1}(n-1)-\theta_{2}(n-1))}{2}}, \varphi(n-1)=2*arctan\left( \frac{2r-sin(\theta_{1}(n-1))-sin{(\theta}_{2}(n-1))}{cos(\theta_{1}(n-1))+cos(\theta_{2}(n-1))} \right).$$

Figure 2 in the main text is simulated by this iterative formula. Regarding the selection of B1 and B2: for the simulation of white noise, refer to the subsequent Section B; for the simulation of power-law noise, refer to the subsequent Section D.

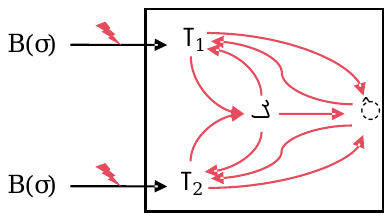

Figure S2 Iterative law diagram

#### B. Approximate solution of simulation 1

Considering Gaussian white noise with noise intensity$\sigma$, it can be written as $\sigma$*B(t), where B(t) ~ N(0,1), N(0,1) represents a Gaussian distribution with a mean of 0 and a variance of 1. According to the properties of variance, there is $\sigma*B(t)\sim N(0,\sigma^{2})$. Since Gaussian white noise does not have temporal correlation, we can simply remember the Gaussian white noise with noise intensity$\sigma$ as $B(\sigma)$, which is a Gaussian white noise with a mean of 0 and a standard deviation of $\sigma$.

Taking $B_{1}$ and $B_{2}$ as $B(\sigma)$, with random initialization for $\theta_{1},\theta_{2}$, and iterating 2500 times with $w_{1}=2.5,w_{2}=4.5$, we analyze the impact of increasing the standard deviation $\sigma$ of Gaussian white noise across various K values. After averaging 100 numerical simulations, we observe that R shows a pattern of initial increase followed by a decrease, as shown in Figure 2 a-d. The following is a further analysis of the influence of each parameter in the model on the coupled-oscillator subsystem.

(1) **Effect of number of iterations T in the model**

By incrementally adjusting the noise intensity from 0 to 4 with a step size of 0.01, we examined the changes in the average level of synchronization, represented by R, throughout the iterative process. The results indicate that the average synchronization R tends to first increase and then decrease as the noise intensity rises with the progression of iterations.

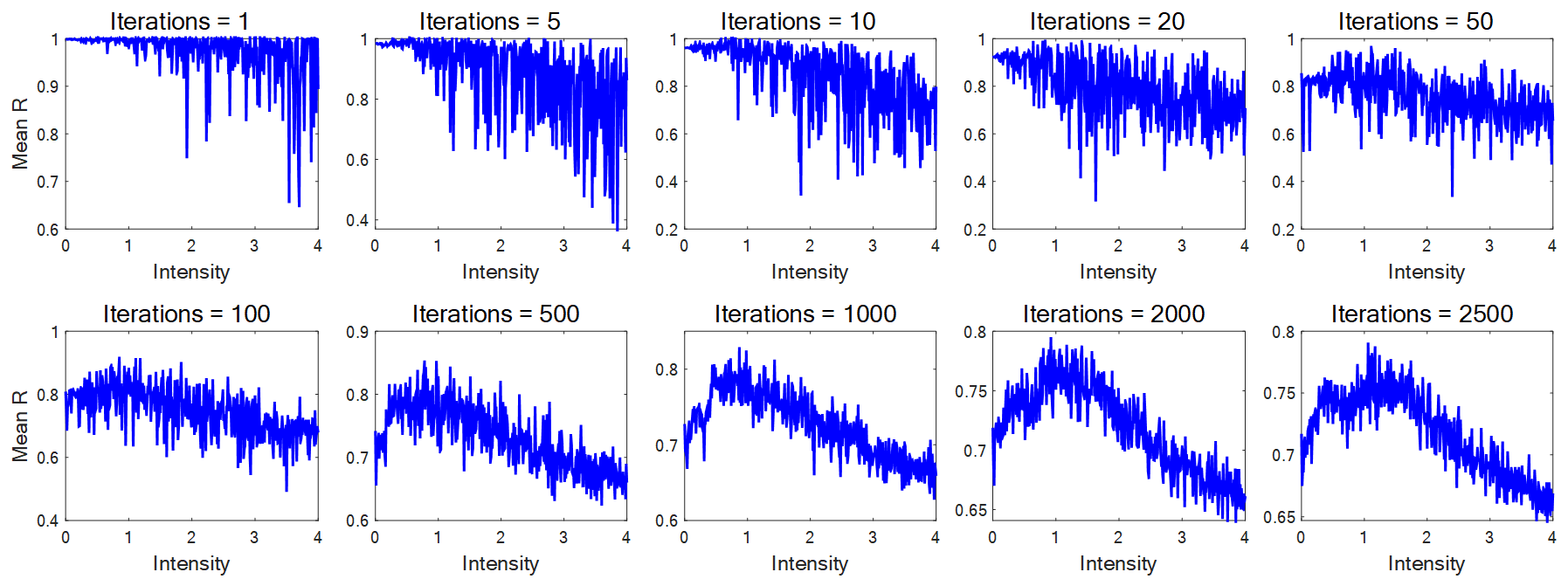

Figure S3 The effect of the number of iterations T on the impact of noise intensity on synchronization R. Parameter setting: iteration step size $dt=0.05$; natural frequency $w_{1}=2,w_{2}=5$; coupling parameter $K=\Delta w=3$.

(2) **Effect of natural frequency** $w$ **in the model**

In the context of a coupled-oscillator system, by subtracting the two equations in the Kuramoto model, we obtain the following differential equation:$d\Delta\theta/dt= \Delta w-\left( K\sin\Delta\theta\right)/2+\Delta B$. Utilizing this equation, it can be deduced that the difference $\Delta w=w_{1}-w_{2}$ between the natural frequencies $w_{1}$ and $w_{2}$ influences $\Delta\theta$. Subsequently, $\Delta\theta$ affects the synchronization R through the formula $R=\sqrt{(1+cos\Delta\theta)/2}$. Without loss of generality, we can set $w_{2}=0$ and vary $w_{1}$ from 0 to 10 to observe the trend of synchronization variation as the noise intensity increases.

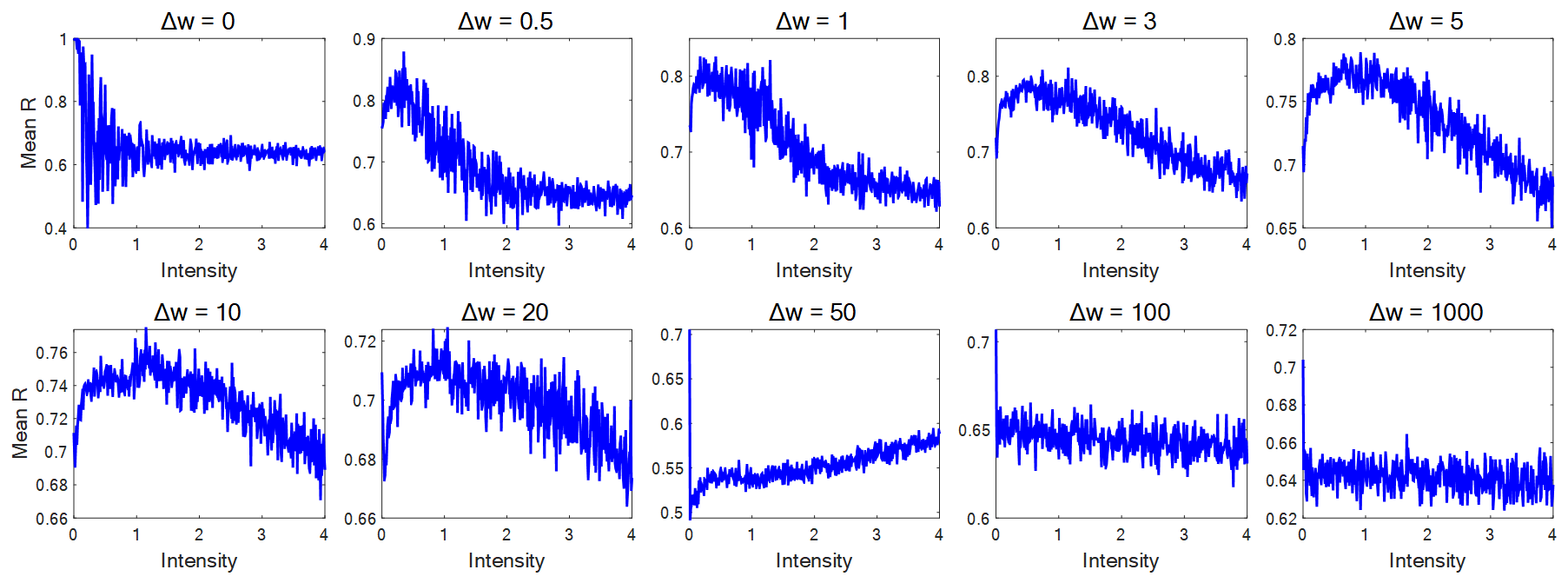

Figure S4 The effect of the natural frequency difference $\Delta w$on the impact of noise intensity on synchronization R. Parameter setting: iteration step size$dt=0.05$; iterations $T=2000$; coupling parameter $K=\Delta w$.

(3) **Effect of coupling parameter** $\mathbf{B}$ **in the model**

According to the literature^[4]^, the critical point for the coupling parameter in the coupled-oscillator model occurs at K=w. Analyzing the simulation results presented in the following figure, a reverse U-shaped trend is observed near the critical point, which is particularly pronounced around K=5.

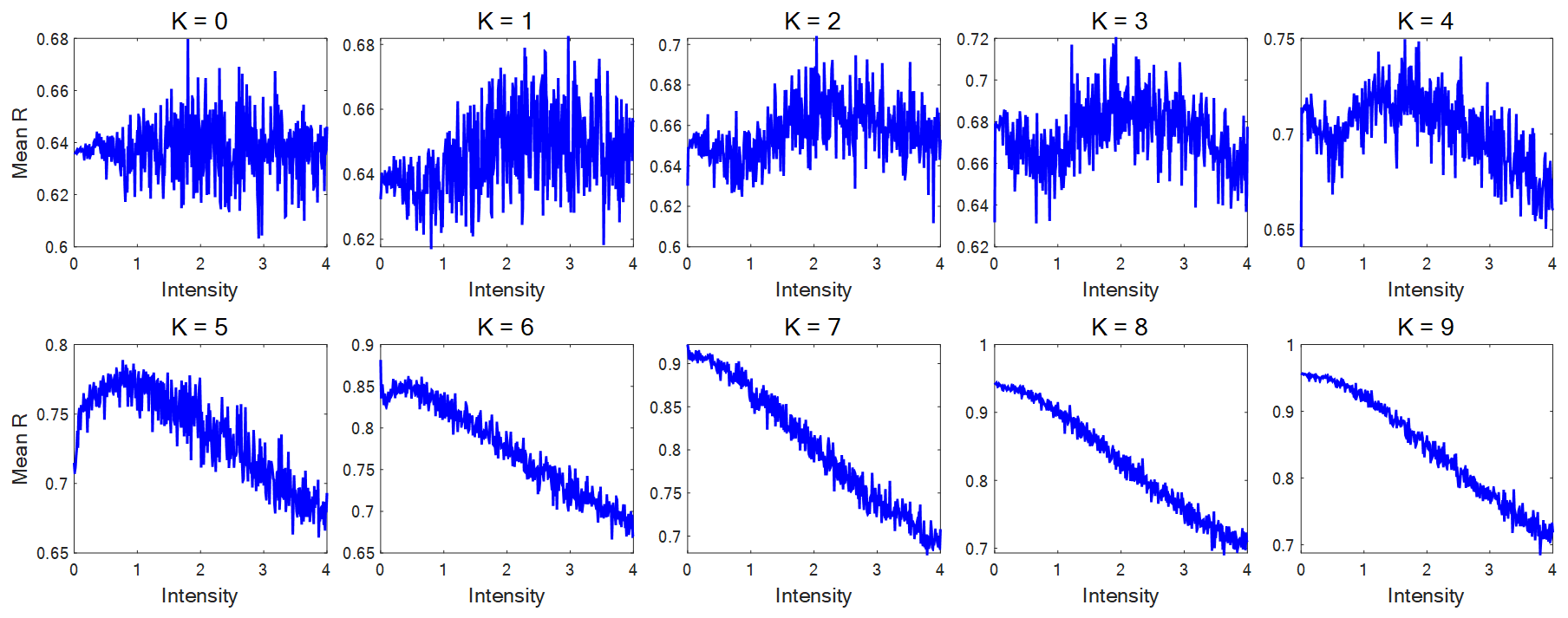

Figure S5 The effect of the coupling parameter K on the impact of noise intensity on synchronization R. Parameter setting: iteration step size$dt=0.05$; iterations $T=2000$; natural frequency$w_{1}=5,w_{2}=0$.

(4) **Effect of iteration step size** $\mathbf{dt}$ **in the model**

Fixing the total length of the iteration at 100 seconds and selecting various iteration step sizes dt ranging from [0.01, 0.05, 0.1, 0.5, 1, 3, 5, 10, 20, 100], we examine the impact of the iteration step size on the synchronization R under the influence of noise B. It is observed that within a smaller range of dt (as shown in the figure below where dt≤0.1), there is an almost uniform pattern (an inverse U-shaped trend), indicating that when considering an interval of 100 seconds, an iteration step size dt≤0.1 may have already reached a state of convergence. In contrast, the second row of the figure with dt values of [0.5, 1, 3, 5, 10, 20, 100] has not yet reached a state of convergence, and thus the results lack reference significance.

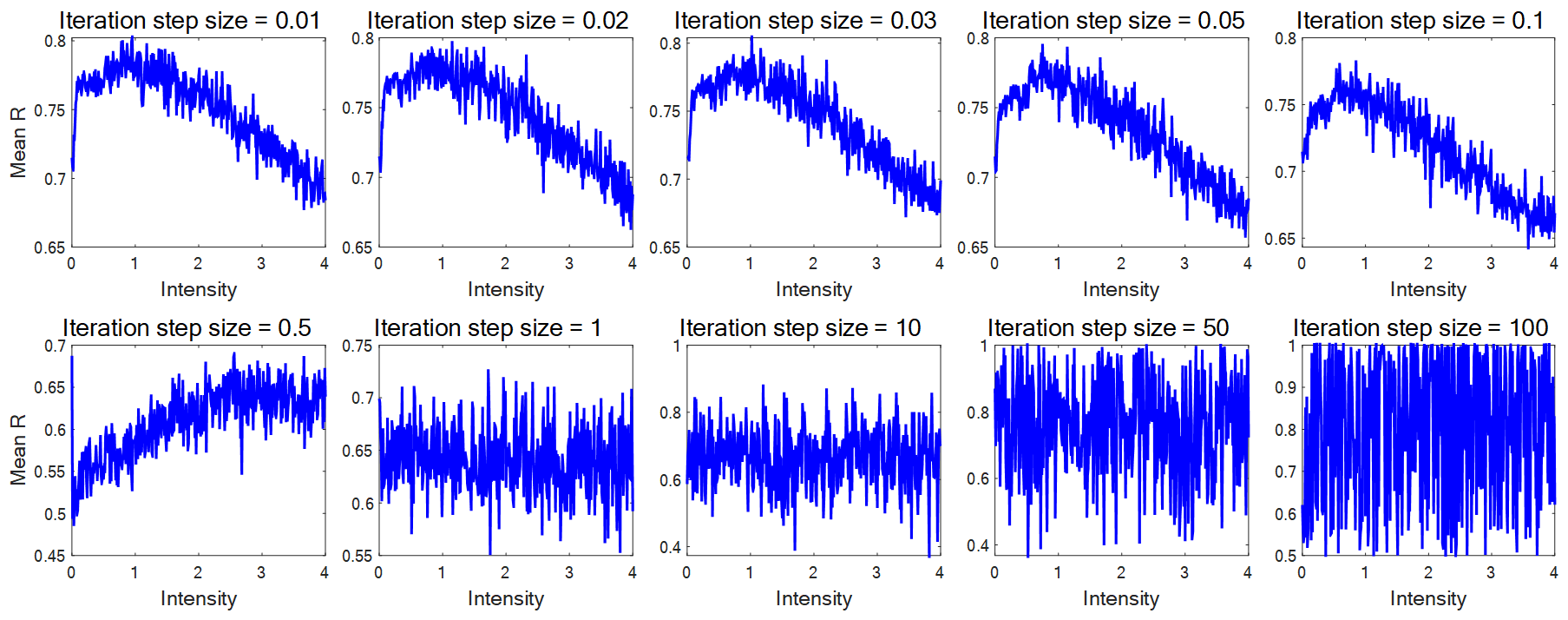

Figure S6 The effect of the iteration step size dt on the impact of noise intensity on synchronization R. Pparameter setting: iterations $T=100/dt$; natural frequency$w_{1}=5,w_{2}=0$; coupling parameter$K=\Delta w=5$.

(5) **How does the intensity of white Gaussian noise affect the synchronization R**

From the previous derivation, $R=\sqrt{(1+cos\Delta\theta)/2}$ shows that R can be determined only by $\Delta\theta$. Combined with the formula $d\Delta\theta/dt= \Delta w-\left( K\sin\Delta\theta\right)/2+\Delta B$, it can be considered that the intensity of noise B affects $\Delta\theta$ through $\Delta B$, and further determines R. The relationship between the three is shown in the figure below.

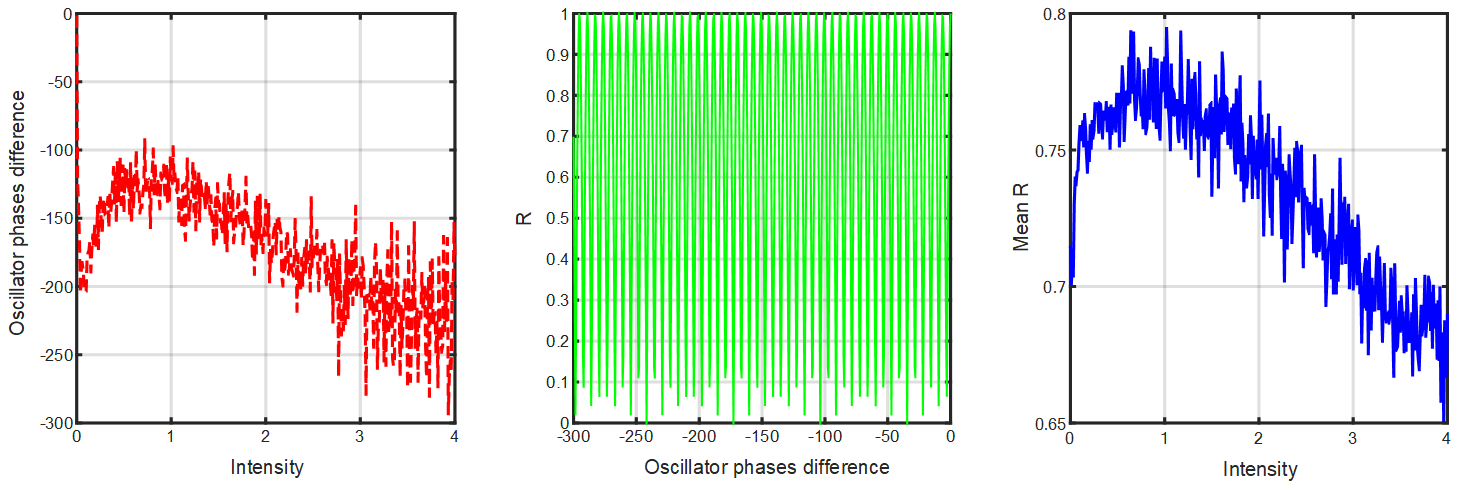

Figure S7 Noise intensity determines synchronization R by affecting the phase difference of the two oscillators. Parameter setting: $K=\Delta w=5$.

#### C. Approximate solution of simulation 2

Using the power-law noise $f(B(\sigma),\beta)$ generated according to the method in Section **I**, by replacing$B_{1}$and$B_{2}$in the iterative formula, we can obtain Simulation 2 in the main text.

Initialize $\theta_{1}$and$\theta_{2}$ randomly, iterate 2500 times, and set $w_{1}=2$and $w_{2}=5$. Under different K values, as the 1/f noise slope β increases, perform numerical simulations 100 times and take the average. It is observed that R exhibits a pattern of initial increase followed by a decrease. This behavior is illustrated in Figures 2 h-k of the main text. Similar to Section B, under the same conditions, we perform an analogous analysis and present the results as follows.

(1) **Effect of number of iterations T in the model**

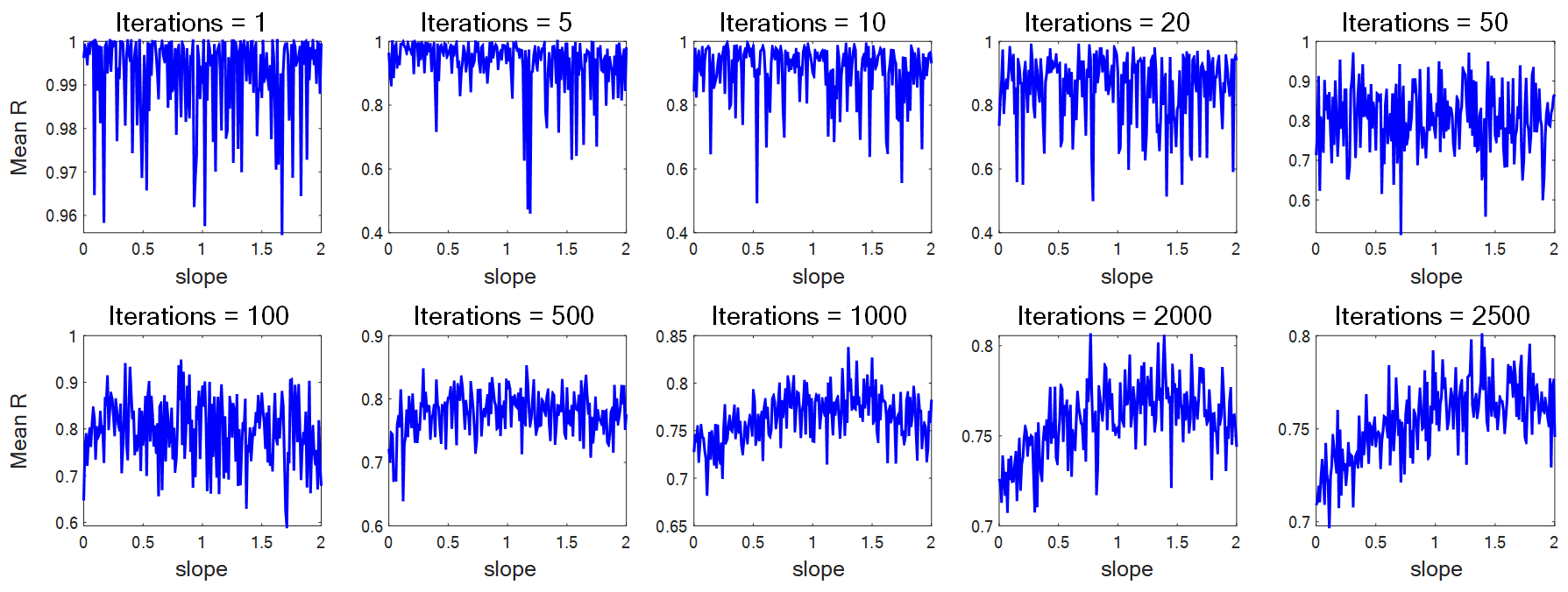

Figure S8 The effect of the number of iterations T on the impact of 1/f slope on synchronization R.

(2) **Effect of natural frequency** $\mathbf{w}$ **in the model**

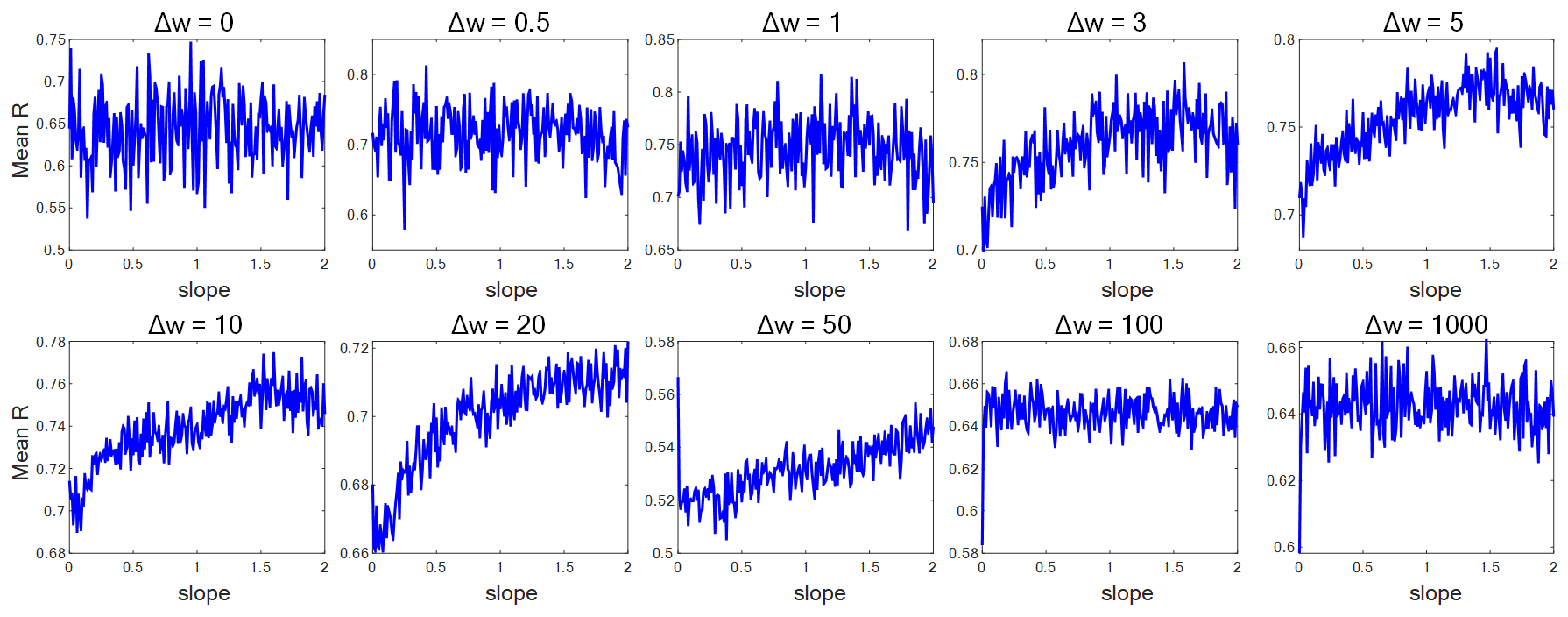

Figure S9 The effect of the natural frequency difference on the impact of 1/f slope on synchronization R.

(3) **Effect of coupling parameter** K **in the model**

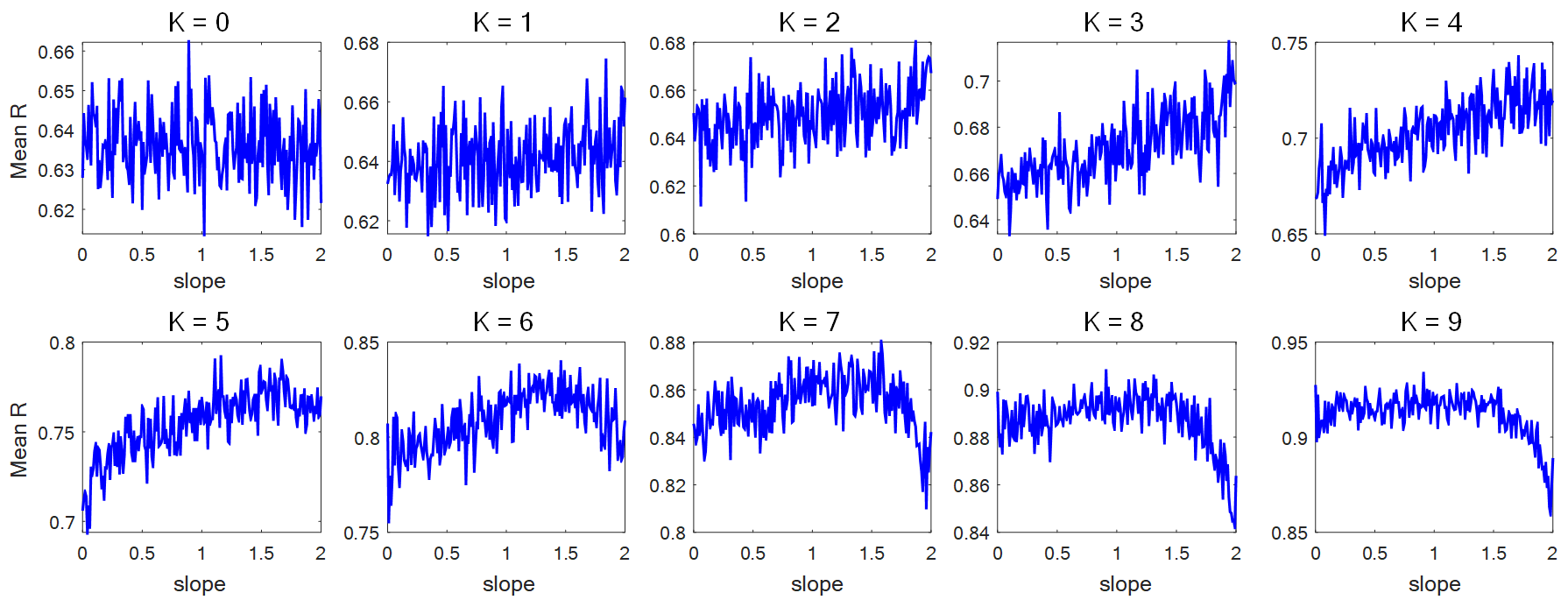

Figure S10 The effect of the coupling parameter K on the impact of 1/f slope on synchronization R.

(4) **Effect of iteration step size** $\mathbf{dt}$ **in the model**

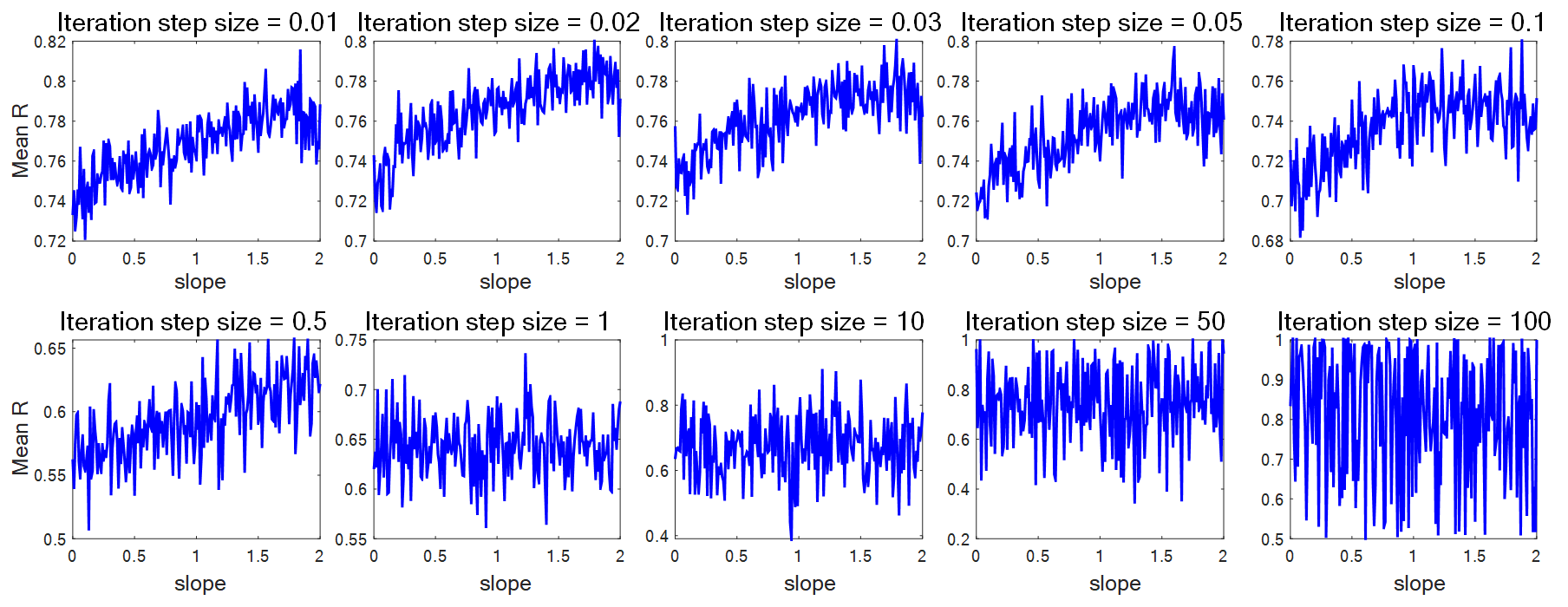

Figure S11 The effect of the iteration step size dt on the impact of noise intensity on synchronization R.

(5) **How does the power-law exponent (slope) of 1/f noise affect the synchronization R**

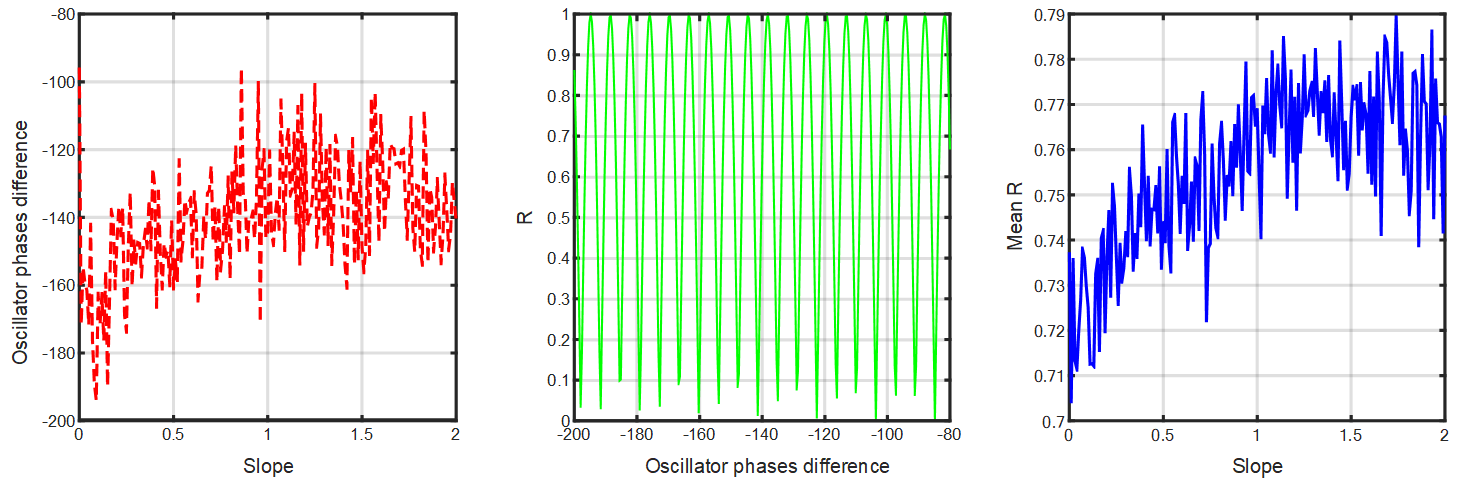

Figure S12 1/f slope determines synchronization R by affecting the phase difference of the two oscillators.

#### D. Analysis by synthesis

(1) **In coupled-oscillator model, the combined effect of intensity and slope on R**

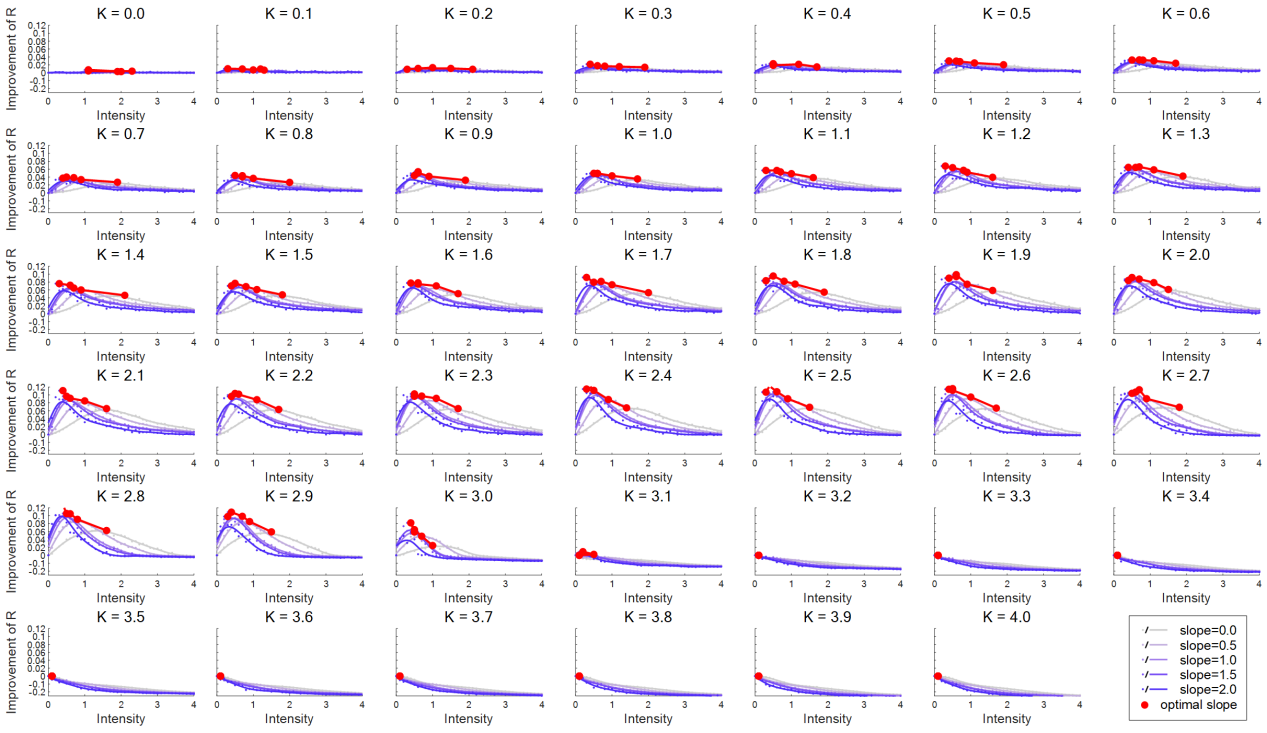

Figure S13 The relationship between the improvement of synchronization R and noise intensity under five different 1/f slopes, with varying coupling parameter K.

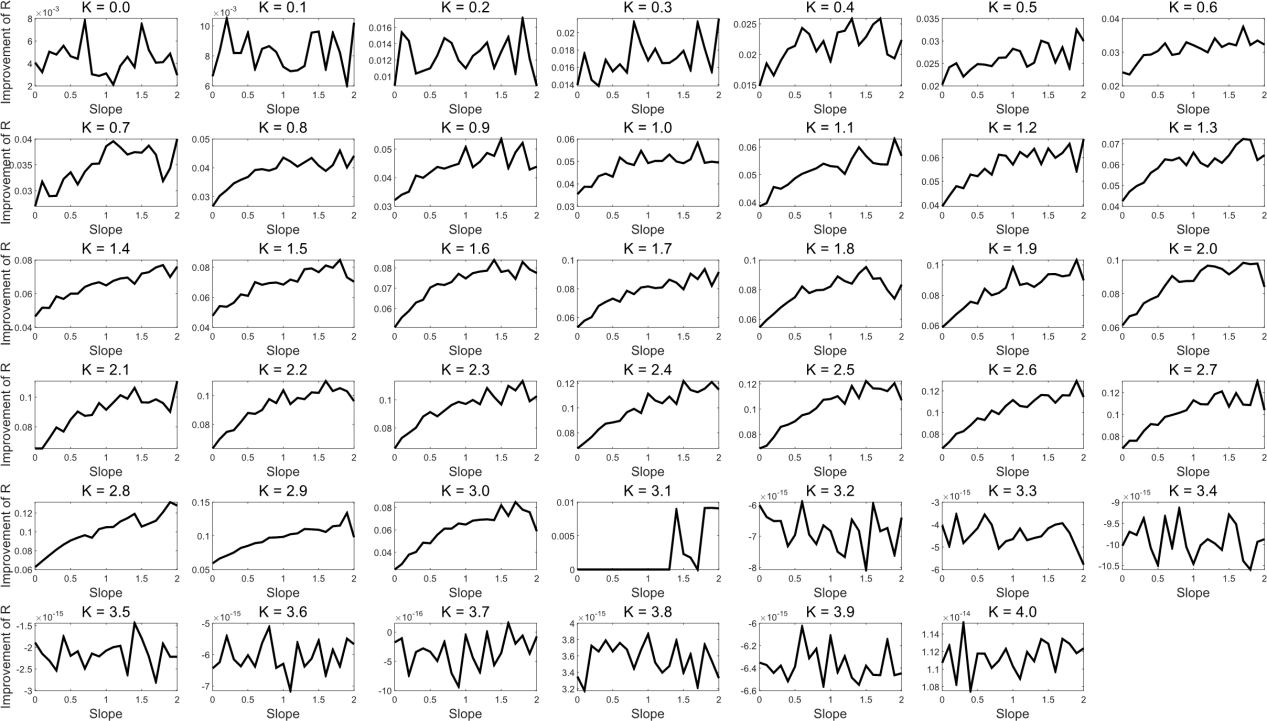

Figure S14 The relationship between the improvement of synchronization R and 1/f slope under different coupling parameter K.

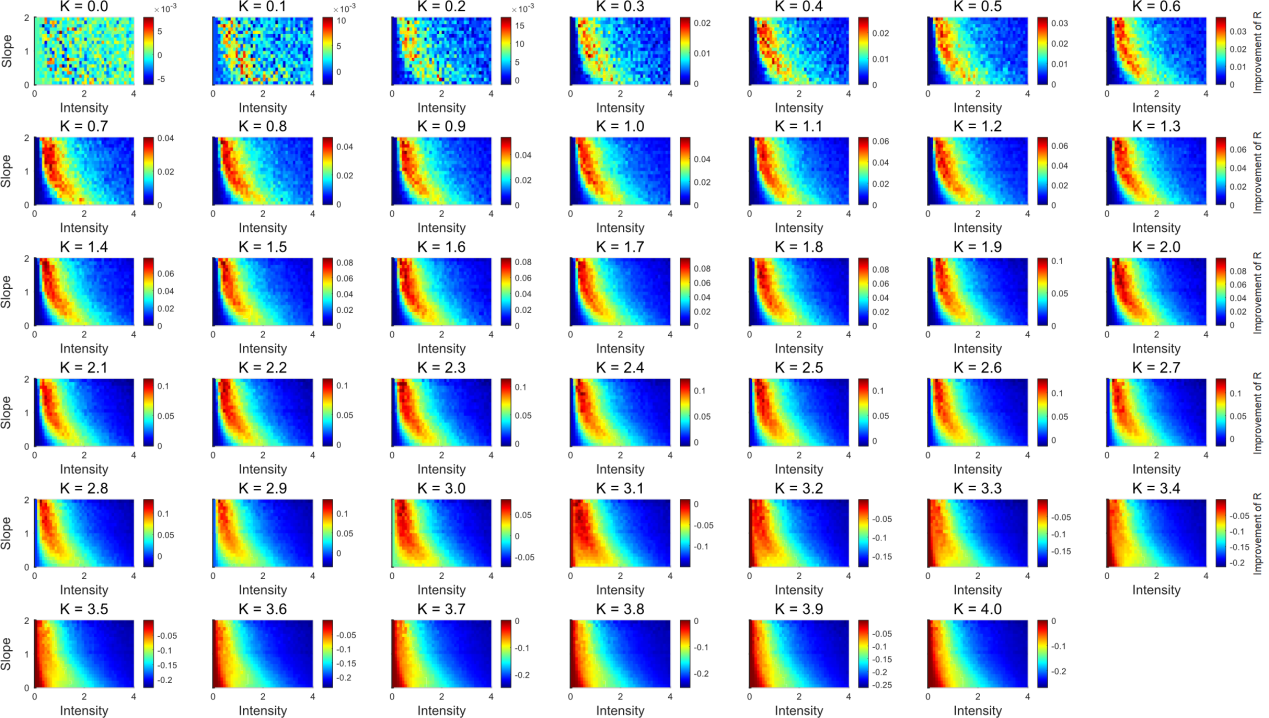

Figure S15 The synergistic effect of noise intensity and noise slope on the enhancement of synchronization R under different coupling parameter K.

Conduct a trend analysis on it (refer to Appendix Part III), and obtain its convexity indicator and growth indicator:

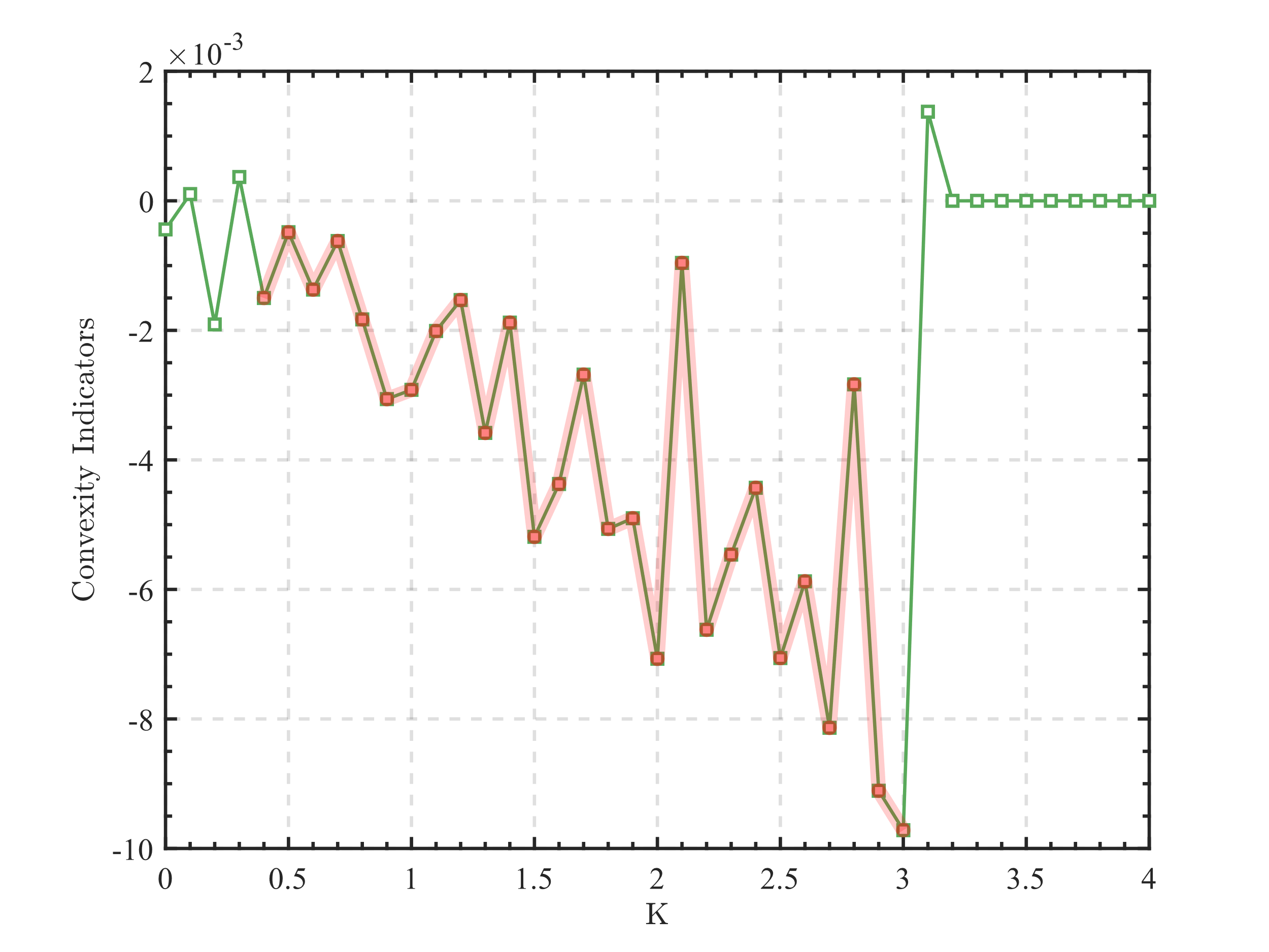

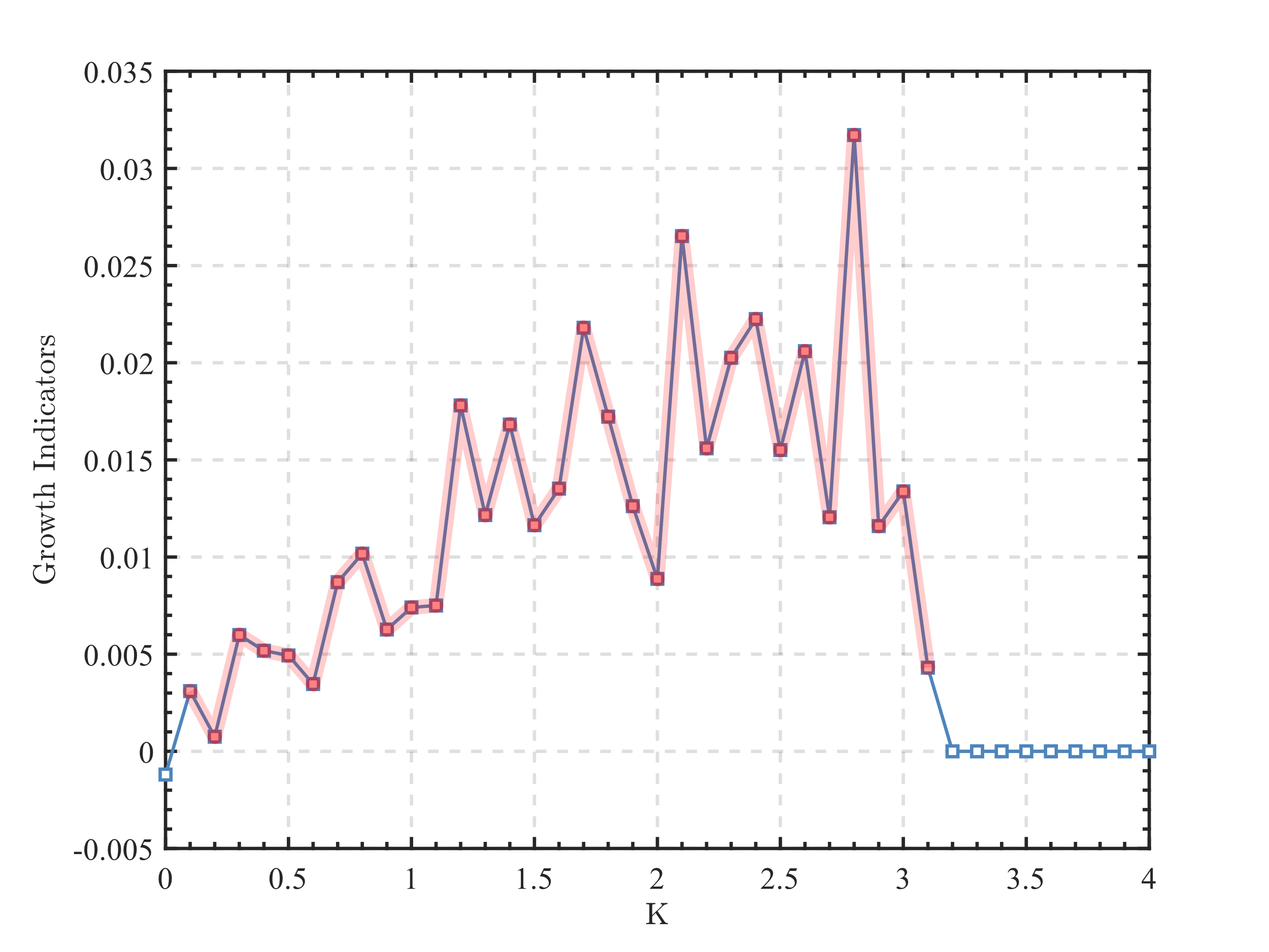

Figure S16 The convexity indicators and growth indicators of the coupled-oscillator model (N=2) under different coupling parameter K.

Accordingly, it is divided into a non-significant impact interval K=[0:0.3], a positive effect interval K=[0.4:3.0], and a negative effect interval K=[3.1:4.0]. Furthermore, the positive effect interval is divided into two segments to observe the changes in its strength. The analysis results after segmentation are presented in Figures 2 e-g of the main text.

**(2) In multi-oscillator model (N=32), the combined effect of intensity and slope on R**

Conduct a trend analysis on it (refer to Appendix Part III), and obtain its convexity indicator and growth indicator:

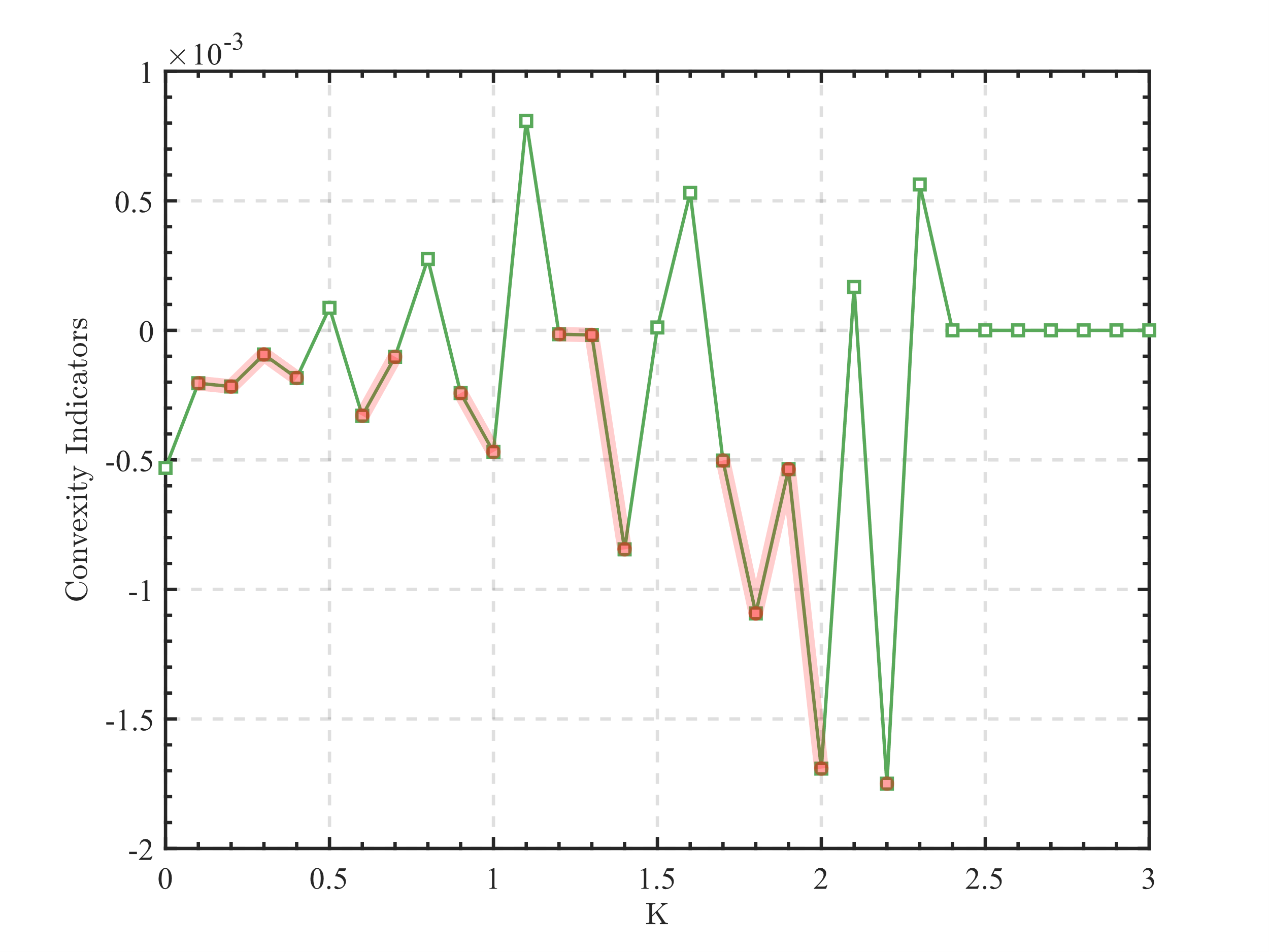

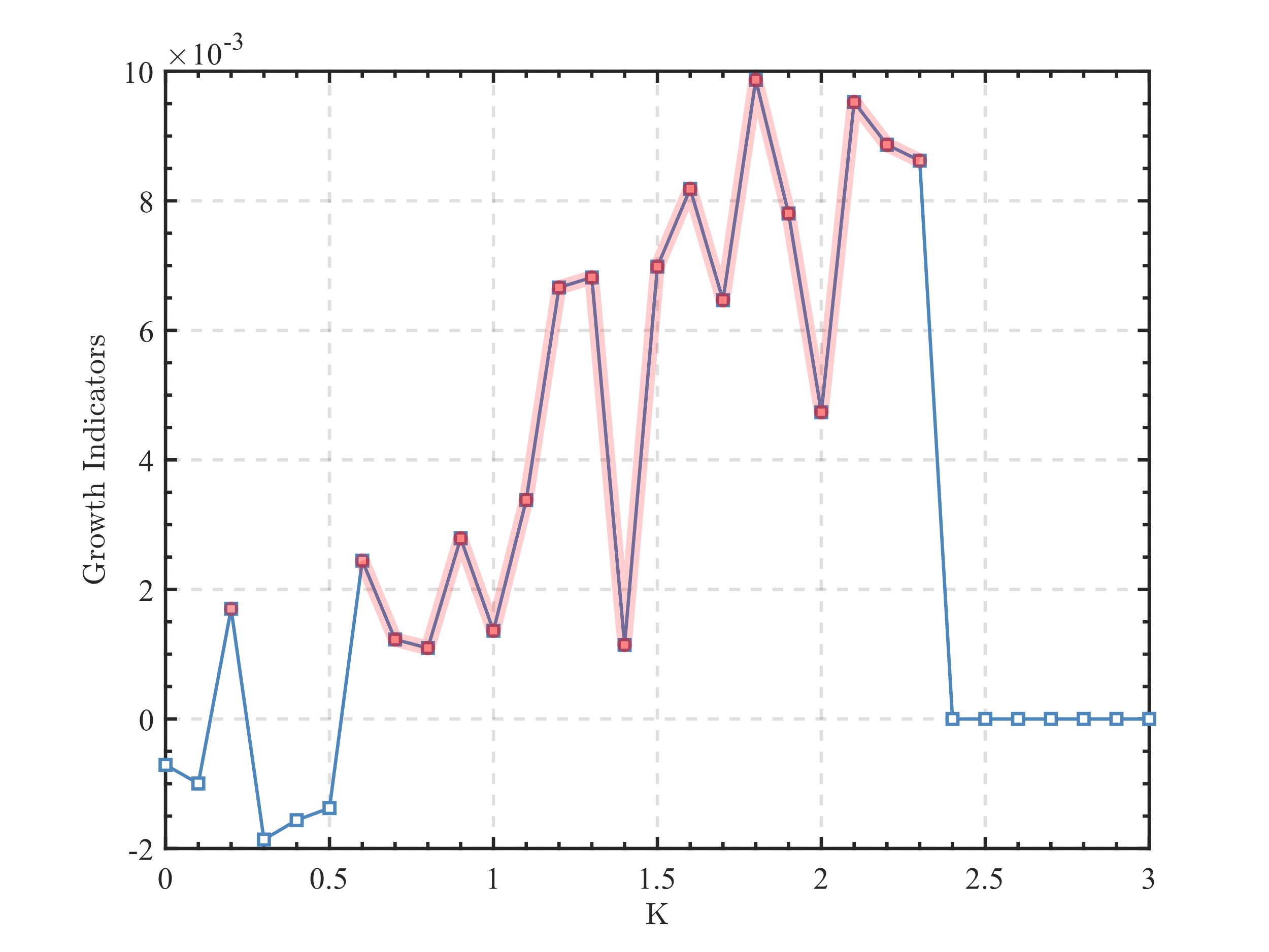

Figure S17 The convexity indicators and growth indicators of the multi-oscillator model (N=32) under different coupling parameter K.

Accordingly, it is divided into four intervals, which are K=[1,6], K=[7,17], K=[18,24], and K=[25,31]. The analysis results after segmentation are shown in Figure S18.

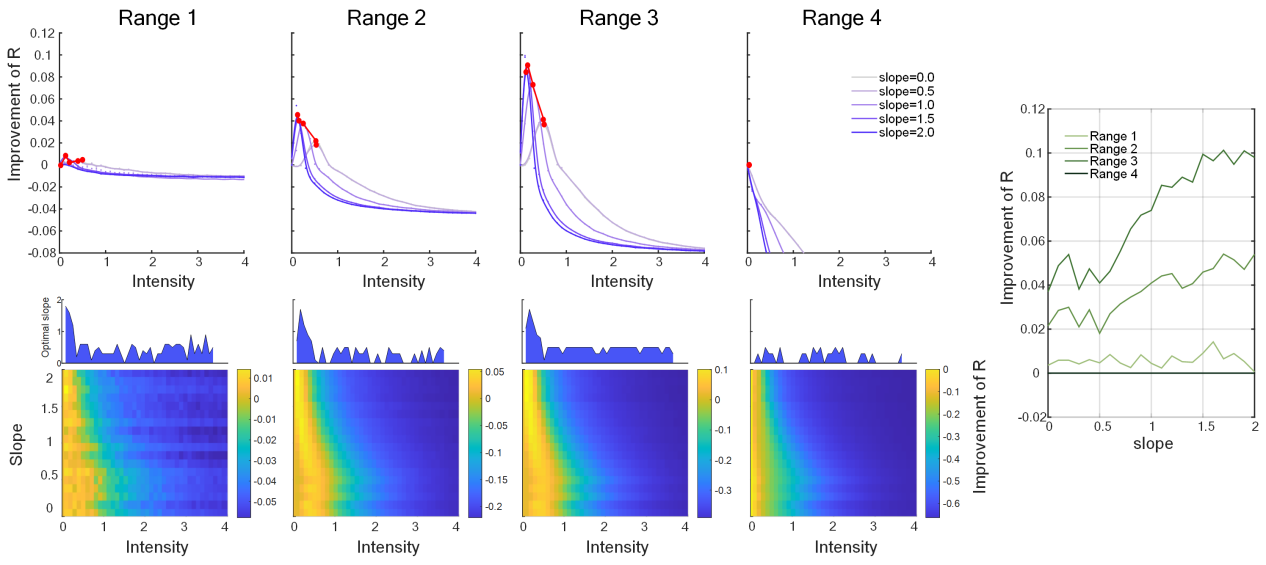

Figure S18 Comprehensive analysis on four coupling parameter intervals.

By comparing the characteristics of the multi-oscillator system (with N=32 oscillators) shown in Supplementary Figure S18 with the coupled-oscillator system (N=2) depicted in Figures 2 e-g of the main text, we observe that both systems follow similar dynamic rules to some extent. Specifically, we reconfirm a common feature: under certain coupling strength conditions, a lower level of noise intensity promotes the synchronization behavior of the system, and an increase in the 1/f slope can further enhance this effect. Notably, compared to the coupled-oscillator model, the multi-oscillator configuration exhibits a distinctive feature: the noise intensity required to achieve optimal synchronization enhancement is significantly reduced. This finding suggests that as the number of oscillators increases, the model's sensitivity to noise and its efficiency in utilizing noise undergo favorable changes for synchronization optimization.

### Ⅲ. Trend quantitative analysis

The human brain is regarded as a highly nonlinear and complex dynamic system, the intrinsic mechanisms of which are difficult to precisely describe and fit using simple mathematical functions. Given that experimental data on human brain activity exhibit discrete characteristics and are often limited in sample size, and considering that many mathematical analysis methods rely on continuous data, while statistical and deep learning approaches require a substantial foundation of data, there are significant limitations in conducting trend analysis, particularly quantitative trend analysis, on experimental data related to human brain activity.

On the other hand, from the perspective of physiological homeostasis, numerous physiological indicators, when responding to various stimuli, typically adhere to the 'optimal moderation' principle. That is, both excessively low and high levels can lead to adverse effects, with the optimal response point often occurring at a moderate level. This results in many experimental data sets exhibiting an inverted U-shaped trend, characterized by an initial rise followed by a decline.

To address this, we creatively proposed a method called trend quantitative analysis, which is composed of three core indicators: Convexity Indicators (CI), Growth Indicators (GI), and Variation Indicators (VI). This method aims to provide a more precise trend analysis for discrete data in experiments. In our study, the trend quantitative analysis is utilized to measure the inverted U-shaped trend, thereby establishing a mutually validating relationship among theory, model, and experimentation.

#### A. Convexity indicators

(1) **Calculation Method**

Considering the case of three points a, b, and c, the vectors $\overset{⃑}{\mathrm{ba}}$ and $\overset{⃑}{\mathrm{bc}}$ are combined with a specific weight $\lambda$ and $\mu$ to form the Convexity Indicator's basic expression $\mathrm{CI}_{0}$. The directionality factor $\delta$ is set based on the relative position of point b to the line segment ac. The formula is as follows:

${\mathrm{CI}_{0}(abc)=\delta\left\| \mu\overset{⃑}{\mathrm{ba}}+\lambda\overset{⃑}{\mathrm{bc}} \right\|}_{2}$	(S7)

where $\delta=\left\{ \begin{aligned} -1, when b is above ac \\ 1, other \end{aligned} \right., \lambda=\frac{|\overset{⃑}{\mathrm{ba}}|}{|\overset{⃑}{\mathrm{ac}}|}, \mu+\lambda=1$. In the subsequent proofs of properties, it is abbreviated as $\mathrm{CI}_{0}$.

Progressing to the case of multiple points, i.e., examining the set $\{P_{0},P_{1}, {\ldots\ldots,P}_{n}\}$, the calculation of the Convexity Indicator is derived from the cumulative and normalized processing of the basic formula involving the first and last points with the intermediate points. This process involves $n+1$ points, with the actual number of summations being $n-1$, yet it is normalized with a denominator of $n\left| P_{0}P_{n} \right|$. This design endows the Convexity Indicator with some desirable mathematical properties. Consequently, for a sequence containing $n+1$ points, the general form of the Convexity Indicator can be expressed as:

$\mathrm{CI}_{n+1}(P_{0}P_{1} {\ldots\ldots P}_{n})=\frac{1}{\mathrm{nL}}\sum_{i=1}^{n-1} \mathrm{CI}_{0}(P_{0}P_{i}P_{n})$	(S8)

where $n+1\geq3$ is the number of points, and $L$ represents the distance between the (n+1)-th point and the starting point along the x-axis. The Convexity Indicator is uniformly denoted as $\mathrm{CI}$ in the main text.

(2) **Properties of Convexity indicators**

**Property 1:** $\frac{\mathbf{2}}{\mathbf{L}_{\mathbf{1}}\mathbf{L}_{\mathbf{2}}}\mathbf{CI}_{\mathbf{0}}$ **is the Taylor expansion approximation of the 2nd derivative. (Approximation of** $\mathbf{CI}_{\mathbf{0}}$**)**

**Proof：**

For any three points $P_{0}(x_{0},y_{0})$,$P_{i}(x_{i},y_{i})$,$P_{n}(x_{n},y_{n})$ in a two-dimensional Cartesian coordinate system, without loss of generality, let $x_{0}$<$x_{i}$<$x_{n}$.Note that the horizontal distance between point $P_{0}$ and point $P_{i}$ is $L_{1}=x_{i}-x_{0}$, the horizontal distance between point $P_{i}$ and point $P_{n}$ is $L_{2}=x_{n}-x_{i}$, and the horizontal distance between point $P_{0}$ and point $P_{n}$ is $L=L_{1}+L_{2}$. Then there is:

$|{\mathrm{CI}_{0}|=\left\| \mu\overset{⃑}{P_{i}P_{0}}+\lambda\overset{⃑}{P_{i}P_{n}} \right\|}_{2}=\left\| \mu(x_{0}-x_{i},y_{0}-y_{i})+\lambda(x_{n}-x_{i},y_{n}-y_{i}) \right\|_{2}$	(S9)

where，

$\lambda=\frac{L_{1}}{L}$ , $\mu=\frac{L_{2}}{L}$	(S10)

Substitute equation (S10) into equation (S9):

$|\mathrm{CI}_{0}|=\sqrt{0^{2}+{(\mu(y_{0}-y_{i})+\lambda(y_{n}-y_{i}))}^{2}}=|\mu(y_{0}-y_{i})+\lambda(y_{n}-y_{i})|.$	(S11)

Since $\lambda L_{2}=\mu L_{1}$，it follows that $\lambda{f'(x_{i})L}_{2}-\mu f'(x_{i})L_{1}=0$，Substituting this into equation (S11)，we get：

$|\mathrm{CI}_{0}|=|\mu(f(x_{0})-f(x_{i})-f'(x_{i})L_{1})+\lambda(f(x_{n})-f(x_{i})+{f'(x_{i})L}_{2})|.$	(S12)

Further, performing a Taylor expansion of $f(x)$ at $x_{i}$ and substituting $x_{0}$ and $x_{n}$ respectively, we have:

$f(x_{0})=f(x_{i})+f'(x_{i})L_{1}+\frac{f''(x_{i})}{2!}{L_{1}}^{2}+R_{i}(L_{1})$	(S13)

$f(x_{n})=f(x_{i})-f'(x_{i})L_{2}+\frac{f''(x_{i})}{2!}{L_{2}}^{2}+R_{i}(L_{2})$	(S14)

Substituting equation (S13) (S14) into equation (S12) , we obtain：

$|\mathrm{CI}_{0}|=|\frac{L_{1}L_{2}}{2}f''(x_{i})+R(L_{1},L_{2})|$	(S15)

Additionally, based on the mathematical significance of $\delta$, it is known that the sign of $\delta$ corresponds to the concavity or convexity determined by the three points. Therefore, when $L$ is sufficiently small or there is no change in concavity or convexity within the $L$ interval, we have:

$\mathrm{CI}_{0}=\frac{L_{1}L_{2}}{2}f''(x_{i})+R(L_{1},L_{2}).$	(S16)

**Property 2: As L approaches 0,** $\frac{\mathbf{2}}{\mathbf{L}_{\mathbf{1}}\mathbf{L}_{\mathbf{2}}}\mathbf{CI}_{\mathbf{0}}$ **converges to the 2nd derivative. (Convergence of** $\mathbf{CI}_{\mathbf{0}}$**)**

**Proof：**

The Lagrange remainders of the Taylor formula in equations (S13) and (S14) are respectively:

$R_{2}(L_{1})=\frac{f'''(\xi_{1})}{3!}{L_{1}}^{3},\xi_{1}\in[x_{0},x_{i}]$	(S17)

$R_{2}(L_{2})=\frac{f'''(\xi_{2})}{3!}{L_{2}}^{3},\xi_{2}\in[x_{i},x_{n}]$	(S18)

Thus, the $R(L_{1},L_{2})$ in equation (S16) can be expressed as:

$R(L_{1},L_{2})=\frac{L_{1}L_{2}}{6(L_{1}+L_{2})}\left( f'''(\xi_{1}){L_{1}}^{2}+f'''(\xi_{2}){L_{2}}^{2} \right), \xi_{1}\in[x_{0},x_{i}],\xi_{2}\in[x_{i},x_{n}]$	(S19)

Substituting into equation (S16), we have:

$\mathrm{CI}_{0}=\frac{L_{1}L_{2}}{2}f''(x_{i})+\frac{L_{1}L_{2}}{6(L_{1}+L_{2})}\left( f'''(\xi_{1}){L_{1}}^{2}+f'''(\xi_{2}){L_{2}}^{2} \right)$	(S20)

As L approaches 0, both $L_{1}$ and $L_{2}$ end to 0, hence:

$\lim_{L\longrightarrow0} \frac{2}{L_{1}L_{2}}\mathrm{CI}_{0}=\lim_{L\longrightarrow0} \left( f''(x_{i})+\frac{L_{1}L_{2}}{6(L_{1}+L_{2})}\left( f'''(\xi_{1}){L_{1}}^{2}+f'''(\xi_{2}){L_{2}}^{2} \right) \right)=f''(x_{i}).$	(S21)

**Property 3:** $\mathbf{CI}_{\mathbf{0}}$ **satisfies the collinearity theorem of plane vectors. (Geometric meaning 1 of** $\mathbf{CI}_{\mathbf{0}}$**)**

Let $P_{c}$ be the endpoint of the vector corresponding to $\mathrm{CI}_{0}$, and let $P_{0}$ and $P_{n}$ be the two points where the entire set of data reaches its extreme horizontal coordinates, then it holds that:

$$\overset{⃑}{P_{i}P_{c}}=\mu\overset{⃑}{P_{i}P_{n}}+\lambda\overset{⃑}{P_{i}P_{0}} 且 \mu+\lambda=1$$

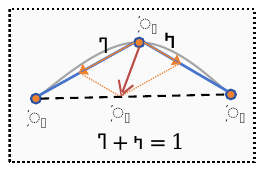

Thus, by the collinearity theorem of plane vectors, $P_{c}$ is collinear with $P_{0}$ and $P_{n}$.

This property indicates that one can find a point $P_{c}$ on the segment $P_{0}P_{n}$ such that Convexity indicators =$|P_{i}P_{c}\boldsymbol{|}$.

**Property 4:** $\mathbf{|}\mathbf{CI}_{\mathbf{0}}\boldsymbol{| \times L =2\times}\mathbf{S}_{\boldsymbol{\Delta}\mathbf{P}_{\mathbf{0}}\mathbf{P}_{\mathbf{i}}\mathbf{P}_{\mathbf{n}}}$**, that is,**$\boldsymbol{cos\alpha= sin\beta}$**. (Geometric meaning 2 of** $\mathbf{CI}_{\mathbf{0}}$**)**

**Proof:**

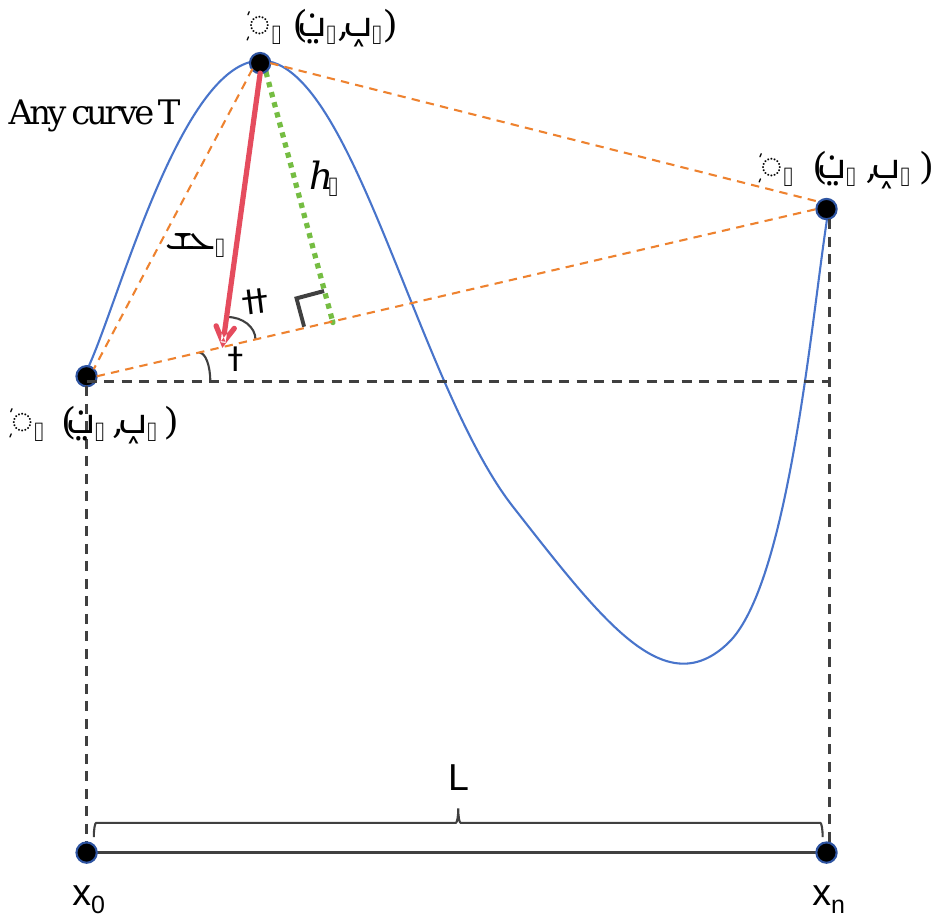

Using the distance from a point to a line formula, we get:

$h_{i}=\frac{\left| (y_{n}-y_{0})x_{i}+(x_{0}-x_{n})y_{i}+(x_{n}y_{0}-x_{0}y_{n}) \right|}{|P_{0}P_{n}|}$	(S22)

According to equation (S11), we obtain:

$\left| {CI}_{0} \right|=\left\| \mu(x_{0}-x_{i},y_{0}-y_{i})+\lambda(x_{n}-x_{i},y_{n}-y_{i}) \right\|_{2}=\sqrt{{(\mu x_{0}+\lambda x_{n}-x_{i})}^{2}+{(\mu y_{0}+\lambda y_{n}-y_{i})}^{2}}$	(S23)

Substituting into equation (S10) and simplifying, we get:

$\left| {CI}_{0} \right|=\left| \frac{(x_{n}-x_{i})y_{0}+(x_{i}-x_{0})y_{n}}{x_{n}-x_{0}}-y_{i} \right|=\frac{\left| (y_{n}-y_{0})x_{i}+(x_{0}-x_{n})y_{i}+(x_{n}y_{0}-x_{0}y_{n}) \right|}{L}$	(S24)

Summarizing the above, we have$\left| {CI}_{0} \right| \times L =h\times|P_{0}P_{n}|$, that is:

$\left| {CI}_{0} \right| \times L =2\times S_{\Delta P_{0}P_{i}P_{n}}$,	(S25)

$\cos\alpha=\sin\beta.$	(S26)

**Property 5:** $\mathbf{|}\mathbf{CI}_{\mathbf{n+1}}\boldsymbol{| \times}\mathbf{L}^{\mathbf{2}}\boldsymbol{\approx}\mathbf{S}_{\boldsymbol{\Delta T-|}\mathbf{P}_{\mathbf{0}}\mathbf{P}_{\mathbf{n}}\mathbf{|}}$**. (Geometric meaning of** $\mathbf{CI}_{\mathbf{n+1}}$**)**

**Proof:**

Substituting Property 4 into equation (S2), we get:

$|\mathrm{CI}_{n+1}(P_{0}P_{1} {\ldots\ldots P}_{n})|=\frac{1}{\mathrm{nL}}\sum_{i=1}^{n-1} |CI(P_{0}P_{i}P_{n})|=\frac{1}{\mathrm{nL}}\sum_{i=1}^{n-1} \frac{2\times S_{\Delta P_{0}P_{i}P_{n}}}{L}=\frac{1}{L^{2}}\sum_{i=1}^{n-1} \frac{|P_{0}P_{n}|}{n}h_{i}$	(S27)

According to the composite trapezoidal rule, we obtain:

$\sum_{i=1}^{n-1} \frac{|P_{0}P_{n}|}{n}h_{i}=\frac{\frac{|P_{0}P_{n}|}{n}}{2}(0+2\sum_{i=1}^{n-1} h_{i}+0)\approx S_{\Delta T-|P_{0}P_{n}|}$	(S28)

Thus, we have:

$|\mathrm{CI}_{n+1}| \times L^{2} \approx S_{\Delta T-|P_{0}P_{n}|}.$	(S29)

where n+1 represents the total number of points, and $S_{\Delta T-|P_{0}P_{n}|}$ represents the area enclosed by the curve T and the line segment $P_{0}P_{n}$.

This property indicates that $|\mathrm{CI}_{n+1}| \times L^{2}$ is a numerical approximation of $S_{\Delta T-|P_{0}P_{n}|}$. Based on the properties of the composite trapezoidal rule, it is known that as $n\longrightarrow\infty$, $|\mathrm{CI}_{n+1}| \times L^{2}$ converges to $S_{\Delta T-|P_{0}P_{n}|}$.

Property 6: The relationship between $\mathbf{CI}_{\mathbf{n}+\mathbf{1}}$ and integration

$$\mathbf{CI}_{\mathbf{n+1}}\boldsymbol{\approx}\frac{\mathbf{1}}{\mathbf{L}^{\mathbf{2}}}\int_{\boldsymbol{x}_{\boldsymbol{0}}}^{\boldsymbol{x}_{\boldsymbol{n}}} \boldsymbol{f(x)dx}\boldsymbol{-}\frac{\boldsymbol{f(}\boldsymbol{x}_{\boldsymbol{0}}\boldsymbol{)+f(}\boldsymbol{x}_{\boldsymbol{n}}\boldsymbol{)}}{\boldsymbol{2}\mathbf{L}}\boldsymbol{.}$$

Proof:

From Property 5, we can deduce:

$|\mathrm{CI}_{n+1}| \times L^{2} \approx S_{\Delta T-|P_{0}P_{n}|}$	(S30)

It is easy to obtain the following geometric relationship:

$S_{\Delta T-|P_{0}P_{n}|}=\left| \int_{x_{0}}^{x_{n}} f(x)dx-\frac{f(x_{0})+f(x_{n})}{2}(x_{n}-x_{0}) \right|$	(S31)

Thus, considering the characteristic of $\delta$, we have:

$\mathrm{CI}_{n+1}\approx\frac{1}{L^{2}}\int_{x_{0}}^{x_{n}} f(x)dx-\frac{f(x_{0})+f(x_{n})}{2L}.$	(S32)

Property 7: The relationship between $\mathbf{CI}_{\mathbf{n}+\mathbf{1}}$ and the 2nd derivative (upper bound estimation)

$\mathrm{CI}_{n+1}\approx\frac{1}{2L^{2}}\int_{x_{0}}^{x_{n}} f''(x)(x-x_{0})(x-x_{n})dx，|\mathrm{CI}_{n+1}| \leq\frac{\mathrm{ML}}{12}+\varepsilon.$	(S33)

Proof:

By performing integration by parts on $\int_{x_{0}}^{x_{n}} f(x)dx$, we obtain:

$\int_{x_{0}}^{x_{n}} f(x)dx-\frac{f(x_{0})+f(x_{n})}{2}(x_{n}-x_{0})=\frac{1}{2}\int_{x_{0}}^{x_{n}} f''(x)(x-x_{0})(x-x_{n})dx$	(S34)

Substituting this into Property 6, we get:

$\mathrm{CI}_{n+1} \times L^{2} \approx\frac{1}{2}\int_{x_{0}}^{x_{n}} f''(x)(x-x_{0})(x-x_{n})dx$	(S35)

Furthermore, if M is the maximum absolute value of the second derivative $f''(x)$ on the interval [$x_{0},x_{n}$], that is:

$\left| f''(x) \right|\leq M$	(S36)

Substituting into equation (S35), we obtain:

$|\mathrm{CI}_{n+1}| \times L^{2} \approx\left| \frac{1}{2}\int_{x_{0}}^{x_{n}} f''(x)(x-x_{0})(x-x_{n})dx \right|\leq\frac{M}{12}{(x_{n}-x_{0})}^{3}=\frac{M}{12}L^{3}$	(S37)

That is, there exists a sufficiently small $\varepsilon>0$, such that when $n\to\infty$, it holds:

$|\mathrm{CI}_{n+1}| \leq\frac{\mathrm{ML}}{12}+\varepsilon.$	(S38)

(3) **Significance of Convexity indicators**

From the core formula S7 of the Convexity Indicator, it can be seen that this indicator quantifies convexity by using the magnitude of the sum of vectors. It transforms complex geometric problems into vector operation problems, which has good operability and provides convenience for further analysis and research. Property 1 and Property 2 establish the relationship between the Convexity Indicator and the second-order derivative, and analyze the Lagrange remainder, obtaining the convergence of CI and the accuracy of the error. Property 3, based on the theorem of collinearity of three points in plane vectors, concretizes the abstractness of concavity and convexity. This geometric construction not only has a clear physical meaning in mathematics but also provides an intuitive geometric interpretation in practical applications. It can also be used to determine the range of CI and further other relationships. The use of this indicator in practical applications may establish a relationship between convexity and specific geometric structures. Properties 4 and 5 provide the relationship between the indicator and the area. In specific experiments, data points often have physiological significance, and when plotted on specific coordinate axes, the area relationship can often represent some physiological meanings, and even correspond to a specific physiological indicator. Properties 6 and 7 respectively provide the relationship between the Convexity Indicator and integration and derivatives, enabling the indicator to be well integrated into the mathematical system for analysis and further extension. In addition, the relationship between the Convexity Indicator and the derivative also provides an upper bound estimate for the indicator.

#### B. Growth indicators and Variation Indicator

In various data analysis inquiries, particularly when dealing with data sequences that exhibit concave or convex characteristics, there is often a keen interest in the extreme points they contain. These extreme points are frequently associated with critical physiological thresholds or pivotal transitional states of systemic mechanisms, which are essential for understanding the regulatory processes of physiology and deciphering the underlying mechanisms. Additionally, the increasing or decreasing trends of data also warrant special attention. Therefore, while conducting an in-depth analysis of the data's concavity or convexity, we typically introduce Growth indicators to analyze variability and hope to detect the presence and characteristics of extreme points.

Growth indicators utilize the traditional principle of approximating the first-order derivative value and employ a composite difference method: initially, calculate the forward difference at the starting point, then successively compute the central difference at the intermediate points, and finally calculate the backward difference at the endpoint. This series of operations collectively constructs an approximation estimate of the first-order derivative that is sensitive to the trends of data changes.

Let there be n+1 data, denoated as $\{P_{0}\left( x_{0},y_{0} \right),P_{1}\left( x_{1},y_{1} \right), {\ldots\ldots,P}_{n}\left( x_{n},y_{n} \right)\}$，, then the calculation formula is as follows:

$Growth indicators=g[x_{0},x_{1}]+\sum_{i=1}^{n-1} g[x_{i-1},x_{i+1}]+g[x_{n-1},x_{n}],$	(S39)

where, $g[x_{i},x_{j}]=\frac{g(x_{j})-g(x_{i})}{x_{j}-x_{i}}=\frac{y_{i}-y_{i}}{x_{j}-x_{i}}$, represents the first-order difference quotient of g(x) at node $x_{i},x_{j}$.

However, real-world data often do not exhibit simple monotonicity. The combination of increasing/decreasing properties and concavity/convexity may not fully capture the trends of the data, especially when focusing on extreme points. According to the theorem of existence of zeros, we can easily understand that each change in sign within the growth indicator often indicates the presence of an extreme value. Therefore, by recording the number of sign changes and directly identifying instances where the difference is zero, we can measure the shifts in data trends.

$Variation Indicator=I\left( g[x_{0},x_{1}]\cdot g[x_{0},x_{2}]\leq0 \right)+\sum_{i=1}^{n-2} I\left( g[x_{i-1},x_{i+1}]\cdot g[x_{i},x_{i+2}]\leq0 \right)+I\left( g[x_{n},x_{n-1}]\cdot g[x_{n},x_{n-1}]\leq0 \right)$	(S40)

where,$I\left( \mathrm{condition} \right)=\left\{ \begin{aligned} 1,if the condition is true \\ 0, if the condition is false \end{aligned} \right.$ is a logical function.

#### C. The application of trend quantitative analysis

In practical applications, Growth indicators and Variation indicators can effectively assist Convexity indicators in analyzing data trends. By analyzing all three together, it is possible to effectively identify and interpret potential turning points and significant trends within a data sequence. Therefore, our trend quantitative indicator can be formulated as:

$TQI=f(CI,GI,VI).$	(S41)

This indicator can be designed with a function $f$ to measure specific trends according to the different problems faced by researchers.

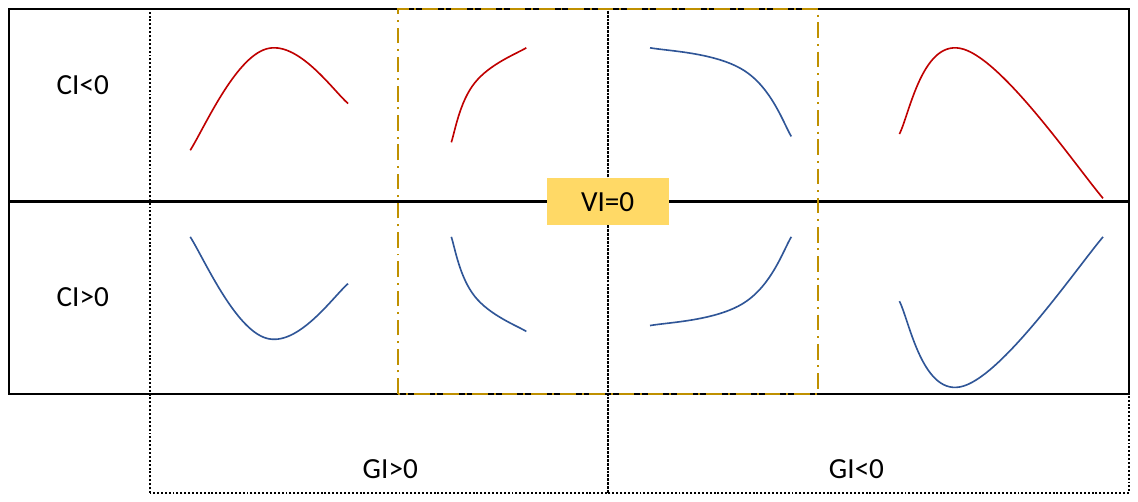

Figure S19 Trend Quantification analysis Selection Trend Illustration.

In the experiments of this study, considering that our experimental scope may not reach the extreme point of the inverted U-shaped curve for everyone, we used: f(CI,GI,VI)=(CI<0 remove VI=0 & GI<0) to select the red trend shown in Figure S19. By multiplying CI with a -1 under the condition where VI=0 & GI<0, we perform the "removal" operation. In order to facilitate understanding, we simply call it "convexity indicator" in the body.

(1) **Application of the Three-Point Formula**

The three-point formula was utilized in the experimental data where the slope was varied. For five different levels of RMT, we conducted a trend quantification analysis of p(MEP ≥ 0.05 mV) values for each individual at slope= 0,1,2.

(2) **Application of the Four-Point Formula**

The four-point formula was applied to the experimental data with varying intensity levels. For five different levels of RMT, we conducted a trend quantification analysis of p(MEP ≥ 0.05 mV) values for each individual at intensity= 0,0.5,1,1.5.

(3) **Application of the Multi-Point Formula**

The Multi-Point formula was employed in simulations of the Kuramoto model when both intensity and slope were altered. We conducted a trend quantification analysis of synchronization R under different noise stimulation conditions of intensity ranging from 0 to 4 with increments of 0.1, and slope ranging from 0 to 2 with increments of 0.1, across various coupling parameters K.

### Statistical tables

Table S1. Statistical analysis results of MEP>50uv probability in experiment 1.

| Experiment1  p (MEP>50uv) | df | F | p |
| --- | --- | --- | --- |
| RMT-3% | 3 51 | 0.9615 | 0.4181 |
| RMT-2% | 3 51 | 1.0503 | 0.3784 |
| **RMT-1**% | 3 **51** | **4.9572** | **0.0043** |
| **RMT** | 3 **51** | **4.9417** | **0.0043** |
| RMT+1% | 3 51 | 2.1663 | 0.1034 |

Table S2. Post hoc analysis results of MEP>50uv probability and Statistical analysis results of RMT-Fit in experiment 1.

| Experiment1  p (MEP>50uv) | Pair-test (FDR corrected) | t | p | Cohen’d |
| --- | --- | --- | --- | --- |
| RMT-1% | **Non V.S. 0.5mA** | **-3.015** | **0.0117** | **0.6518** |
|  | **Non V.S. 1mA** | **-3.316** | **0.0098** | **0.8174** |
|  | **Non V.S. 1.5mA** | **-2.713** | **0.0178** | **0.6390** |
|  | 0.5mA V.S. 1mA | -0.301 | 0.7642 | - |
|  | 0.5mA V.S. 1.5mA | 0.301 | 0.7642 | - |
|  | 1mA V.S. 1.5mA | 0.603 | 0. 7642 | - |
| RMT | **Non V.S. 0.5mA** | **-2.498** | **0.0419** | **0.6616** |
|  | **Non V.S. 1mA** | **-3.628** | **0.0038** | **0.9789** |
|  | **Non V.S. 1.5mA** | **-2.379** | **0.0419** | **0.6389** |
|  | 0.5mA V.S. 1mA | -1.130 | 0.3162 | - |
|  | 0.5mA V.S. 1.5mA | 0.119 | 0.9058 | - |
|  | 1mA V.S. 1.5mA | 1.249 | 0.3162 | - |
| Experiment1 | df | F | | p |
| RMT-Fit (3,18,3,4,7) | **42** | **3.173** | | **0.0339** |
|  | Pair-test (FDR corrected) | t | p | Cohen’s d |
|  | Non V.S. white | 1.263 | 0.3196 |  |
|  | **Non V.S. pink** | **2.857** | **0.0387** | **0.641935362041072** |
|  | Non V.S. brown | 2.028 | 0.1455 |  |
|  | white V.S. pink | 1.594 | 0.2360 | - |
|  | white V.S. brown | 0.765 | 0.4485 | - |
|  | pink V.S. brown | -0.829 | 0.4485 | - |

Table S3. Statistical analysis results of MEP>50uv probability in experiment 2.

| Experiment2  p (MEP>50uv) | df | F | p |
| --- | --- | --- | --- |
| RMT-4% | 3 46.256 | 1.5043 | 0.2259 |
| **RMT-2**% | 3 **46.305** | **3.5201** | **0.0222** |
| **RMT** | 3 **46.267** | **5.3549** | **0.0029** |
| **RMT+2**% | 3 **46.049** | **4.1374** | **0.0112** |
| RMT+4% | 3 46.085 | 2.1495 | 0.1069 |

Table S4. Post hoc analysis results of MEP>50uv probability in experiment 2.

| Experiment1  p (MEP>50uv) | Pair-test (FDR corrected) | t | p | Cohen’s d |
| --- | --- | --- | --- | --- |
| RMT-2% | Non V.S. white | -1.061 | 0.3529 | - |
|  | Non V.S. pink | -1.500 | 0.2289 |  |
|  | **Non V.S. brown** | **-3.085** | **0.0200** | **-0.912870929175277** |
|  | white V.S. pink | -0.485 | 0.6297 | - |
|  | white V.S. brown | -2.025 | 0.1451 |  |
|  | pink V.S. brown | -1.453 | 0.2289 | - |
| RMT | **Non V.S. white** | **-2.757** | **0.0163** | **-0.610729415508371** |
|  | **Non V.S. pink** | **-3.041** | **0.0113** | **-0.551113383559555** |
|  | **Non V.S. brown** | **-3.466** | **0.0066** | **-0.930556346764566** |
|  | white V.S. pink | -0.403 | 0.7843 | - |
|  | white V.S. brown | -0.709 | 0.7226 | - |
|  | pink V.S. brown | -0.275 | 0.7843 | - |
| RMT+2% | Non V.S. white | -0.741 | 0.4624 | - |
|  | **Non V.S. pink** | **-3.066** | **0.0211** | **-0.605941537310370** |
|  | Non V.S. brown | -2.222 | 0.0618 | -0.499039810030499 |
|  | white V.S. pink | -2.357 | 0.0618 | -0.550832002957130 |
|  | white V.S. brown | -1.481 | 0.2173 | - |
|  | pink V.S. brown | 0.940 | 0.4221 | - |

Table S5. Statistical analysis and post hoc results of RMT-Fit in experiment 2.

| Experiment2 | df | F | | p |
| --- | --- | --- | --- | --- |
| RMT-Fit (2,15,17,19) | **43.14** | **9.0305** | | **9.316e-05** |
|  | Pair-test (FDR corrected) | t | p | Cohen’s d |
|  | **Non V.S. white** | **2.673** | **0.0207** | **0.659379835936374** |
|  | **Non V.S. pink** | **4.471** | **0.0003** | **0.904893109990299** |
|  | **Non V.S. brown** | **4.129** | **0.0005** | **0.817228915937777** |
|  | white V.S. pink | 1.923 | 0.0909 | 0.503994929987514 |
|  | white V.S. brown | 1.455 | 0.1828 | - |
|  | pink V.S. brown | -0.536 | 0.5944 | - |

Table S6. Statistical analysis results of MEP amplitude in experiment 2.

| Experiment2  p (MEP>50uv) | df | F | p |
| --- | --- | --- | --- |
| RMT-4% | 3 46.245 | 2.0331 | 0.1223 |
| RMT-2% | 3 45.532 | 2.8001 | 0.05052 |
| RMT | 3 46.019 | 7.4502 | 0.0003625 |
| RMT+2% | 3 45.423 | 6.8284 | 0.0006782 |
| RMT+4% | 3 45.797 | 1.8969 | 0.1434 |

Table S7. Statistical analysis results of MEP amplitude in experiment 2.

| Experiment1  p (MEP>50uv) | Pair-test (FDR corrected) | t | p | Cohen’s d |
| --- | --- | --- | --- | --- |
| RMT | Non V.S. white | -1.124 | 0.2666 |  |
|  | **Non V.S. pink** | **-2.300** | **0.0513** | **-0.498989681876708** |
|  | **Non V.S. brown** | **-4.356** | **0.0004** | **-1.27837673662160** |
|  | white V.S. pink | -1.225 | 0.2666 | - |
|  | white V.S. brown | **-3.232** | **0.0066** | -0.679044694713797 |
|  | pink V.S. brown | -1.867 | 0.1018 | - |
| RMT+2% | **Non V.S. white** | **-2.453** | **0.0355** | **-0.540097621939687** |
|  | **Non V.S. pink** | **-4.251** | **0.0006** | **-0.871882986119556** |
|  | **Non V.S. brown** | **-2.934** | **0.0152** | **-0.721705341962714** |
|  | white V.S. pink | -1.905 | 0.0938 | -0.357212714896645 |
|  | white V.S. brown | -0.481 | 0.6327 | - |
|  | pink V.S. brown | 1.445 | 0.1856 | - |

1. The origin of $\sqrt{dt}$ in the formula, please refer to the literature [1][2].
